# Repurposing dopamine D_3_ receptor fluorescent ligands for D_2_ receptor

**DOI:** 10.64898/2026.10.07.756687

**Authors:** Kätlin Ots, Tõnis Laasfeld, Kaspar Hollo, Mihkel Ilisson, Sandra Ortigueira Noya, Fabian Grassl, Dimitra Tousi, Hanna-Riia Allas, Maria Majellaro, Erki Enkvist, Markus Heinrich, Eddy Sotelo, Anni Allikalt

## Abstract

The dopamine D_2_ receptor is a principal target of antipsychotic and antiparkinsonian drugs, yet well-characterized fluorescent ligands for the D_2_ receptor remain scarce compared with its relative, the D_3_ receptor. Here we show that fluorescent ligands developed for the D_3_ subtype can be repurposed as tools for studying the D_2_ receptor pharmacology. Using a machine-learning-assisted, live-cell fluorescence microscopy assay, we screened twelve fluorescent probes and identified eight ligands that bound specifically to the D_2_ receptor at 10 nM. Four ligands exhibited classical saturable binding, yielding K_d_ in the nanomolar range, whereas the remaining probes displayed unsaturated, time-dependent binding profiles. Kinetic analyses revealed substantial differences in association and dissociation behavior, highlighting the importance of time-resolved measurements. Competition binding with dopamine and haloperidol yielded affinity estimates consistent with literature values. These results expand the toolkit of validated D_2_ receptor fluorescent probes and support cross-receptor repurposing as a strategy for fluorescent ligand development.

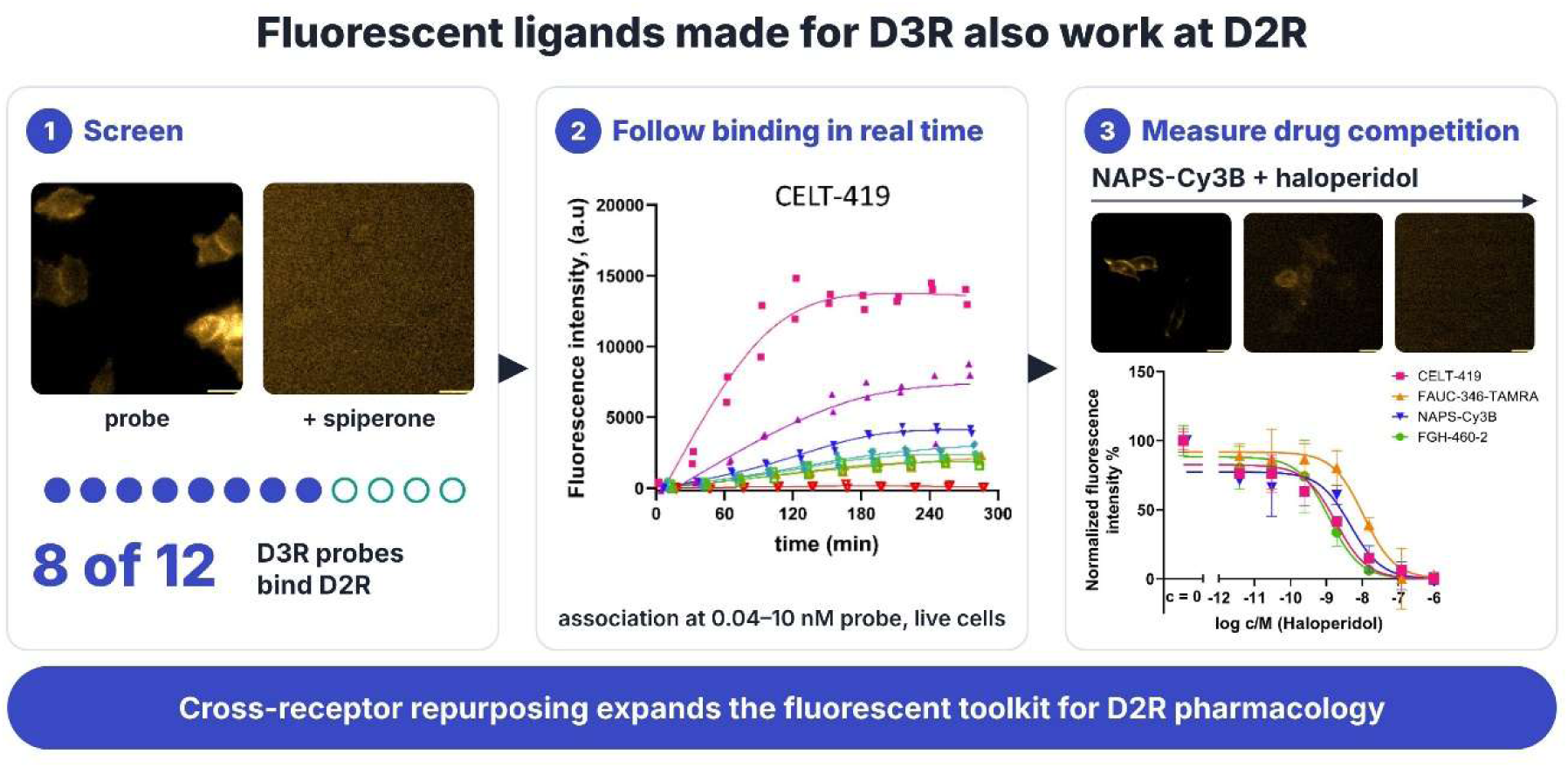

## Introduction

The dopamine D_2_-like receptor family, comprising the D_2_, D_3_, and D_4_ receptor subtypes, is central to dopaminergic neurotransmission^1^ and remains one of the most heavily targeted receptor families in clinical pharmacology. The D_2_ receptor, the most abundant and widely studied member of this family, is expressed throughout the nigrostriatal, mesolimbic, and mesocortical dopaminergic pathways, where it regulates motor output and reward-related behaviour^2^, and is the principal target of essentially all approved antipsychotic drugs^3,4^. It is further implicated in the pathophysiology of schizophrenia, as well as in Parkinson’s disease, where D_2_-directed agonists are used to compensate for dopaminergic neuron loss.^5,6^ The closely related D_3_ receptor has attracted growing interest as a therapeutic target in its own right, owing to its restricted expression in limbic brain regions and its involvement in substance use disorders, impulse control disorders, and as an adjunct target in Parkinson’s disease and schizophrenia pharmacotherapy.^7,8^ Because D_2_ and D_3_ receptors share a high degree of sequence and structural conservation within their orthosteric ligand-binding pocket^4,9–11^, selective and nonselective fluorescent ligands have been developed and characterized for both receptors, enabling their use in a range of fluorescence-based binding assays^12–16^.

Historically, the quantification of ligand-receptor binding has relied heavily on radioligand binding assays^17^, which, despite their sensitivity and long-standing validation, require handling of radioactive material, are typically restricted to membrane preparations or lysed cell suspensions, and provide no spatial or single-cell resolution^18^. Over the past two decades, fluorescence-based binding assays have emerged as a safer, more accessible, and increasingly versatile alternative, exploiting the sensitivity of modern fluorescence detection^16,18–20^ together with the growing chemical toolkit available for conjugating fluorophores to receptor ligands without abolishing their binding activity^12,15,21–23^. Several complementary fluorescence-based formats have since been developed, each offering distinct trade-offs in sensitivity, throughput, and biological context. Fluorescence polarization/anisotropy assays measure the binding-dependent change in the rotational mobility of a fluorescent ligand and are well suited to high-throughput screening in homogeneous, plate-based formats, but typically require relatively high protein concentrations and provide no spatial or single-cell information^20^. Proximity-based methods such as Förster resonance energy transfer (FRET) and nanoluciferase with bioluminescence resonance energy transfer (NanoBRET) offer high sensitivity and are readily adaptable to automated screening, but depend on co-expressing or fusing a second, often genetically engineered, energy donor or acceptor moiety with the receptor, and likewise report only bulk, population-averaged binding^24–27^. Flow cytometry-based assays enable single-cell quantification of fluorescent ligand binding at higher throughput than microscopy, but sacrifice spatial resolution and require cells to be maintained in suspension, limiting their compatibility with adherent cell types and with the study of dynamic subcellular processes^16,18,28^.

Among these approaches, fluorescence microscopy-based binding assays offer a particularly powerful complementary format. Unlike ensemble techniques such as the methods mentioned above, microscopy enables single-cell, spatially resolved visualization of ligand-receptor interactions directly at the plasma membrane, in real time, without requiring receptor purification or homogeneous cell populations^15,16^. This makes microscopy-based assays well suited not only for affinity and kinetic characterization, but also for probing more complex aspects of receptor pharmacology, including internalization, trafficking, and subcellular localization, within a physiologically relevant cellular context^29,30^. This spatial resolution and single-cell sensitivity, however, typically come at the cost of somewhat lower throughput and greater image-analysis complexity relative to plate-based formats^29–31^ ‒ a trade-off that nonetheless makes microscopy well suited to the binding questions addressed in this study.

Despite the pharmacological importance of D_2_, the number of well-characterized fluorescent ligands available for this receptor remains surprisingly limited^12,32–36^. Generating a fluorescent probe that retains sufficient affinity, selectivity, and photophysical quality for robust binding assays typically requires extensive medicinal chemistry optimization, and for D_2_ this effort has yielded only a handful of validated tools relative to the wide range of unlabeled reference antagonists and agonists available. This scarcity constrains the routine implementation of fluorescence-based binding and screening assays for D_2_, and by extension for antipsychotic and antiparkinsonian drug discovery efforts that rely on this receptor. Nevertheless, the abundance of known D_2_ receptor-targeting pharmacophores, together with resolved receptor structures^4,37^ and an expanding repertoire of fluorescent labels, suggests that suitable probes could in principle be readily designed. Moreover, given the conserved orthosteric binding site shared across the D_2_-like receptor family, several fluorescent ligands already developed for the closely related D_3_ receptor represent plausible candidate probes for the D_2_ receptor.

To address this gap, we established a fluorescence microscopy-based binding assay platform capable of quantifying total and nonspecific ligand binding, resolving binding kinetics, and determining equilibrium dissociation constants (K_d_) and competition binding affinities (K_i_) directly in live cells expressing D_2_. The same approach can be used as a starting point for similar panel screens in the future.

Cross-receptor repurposing of this kind has previously proven effective: for example, aminergic GPCR probes have shown useful cross-reactivity across serotonin, dopamine, histamine, and adrenergic receptor subtypes in time-resolved FRET assays^38^ while even a machine-learning based model has been developed that successfully proposes fluorescent ligand structures with promiscuous affinity to many receptor subtypes^39^. This suggests that repurposing may offer a general strategy for rapidly expanding the fluorescent toolkit available for structurally related receptors. In this study, we evaluate the binding of 12 fluorescent ligands at D_2_ using fluorescence microscopy. We chose eight probes with the best signal-to-noise ratio and characterized their saturation binding, association and dissociation kinetics. Then we performed competition binding experiments with four fluorescent ligands using the endogenous agonist dopamine and the antagonist haloperidol. Together, these results demonstrate that D_3_ receptor-targeting fluorescent ligands can be effectively repurposed as D_2_ probes, providing new tools for D_2_ receptor binding assays and establishing a broadly applicable strategy for expanding the fluorescent ligand toolbox within the dopamine receptor family and, potentially, other structurally related GPCR subfamilies.

## Methods

### Cell culture

For microscopy experiments, human embryonic kidney cells (HEK293) expressing the human dopamine D_2L_ receptor (encoded by the gene DRD2) were cultured in BioLite 6-well plates (Thermo Fisher Scientific). The stable cell line expressing the dopamine D_2L_ receptor was developed by Dr. Reet Reinart-Okugbeni (HEK293-D2R)^40^. Cells were maintained in high-glucose (4.5 g/L) Dulbecco’s Modified Eagle Medium (DMEM) (Corning) supplemented with 9% fetal bovine serum (FBS), a mixture of antibiotics (100 U/mL penicillin, 0.1 mg/mL streptomycin, and 0.25 μg/mL amphotericin B) (Sigma-Aldrich), and 400 μg/mL geneticin (G418; MedChemExpress). Cells were cultured in an incubator at 37 °C in a humidified atmosphere containing 5% CO₂. An Olympus CKX31 microscope was used for routine assessment of HEK293-D2R cell confluency, attachment, and morphology. All media and buffer solutions were pre-warmed to 37 °C to avoid exposing the cells to sudden temperature changes. For the assay, cells grown in 6-well cell culture plates (BioLite, Thermo Scientific) were washed with 1 mL of Dulbecco’s phosphate-buffered saline (DPBS, without Ca^2+^ and Mg^2+^; Corning) and incubated with 0.5 mL of 0.05% trypsin solution (Capricorn Scientific) for several minutes at 37 °C in a humidified atmosphere containing 5% CO₂ to detach the cells. Subsequently, 1 mL of DMEM culture medium was added, and the cells were gently detached from the plate surface. The cell suspension was then transferred to a 15 mL tube containing an additional 2 mL of DMEM medium. The cells were centrifuged for 5 minutes at 800 × *g* and the resulting cell pellet was resuspended in 1 mL of culture medium, and cell viability and density were determined by adding a 0.2% trypan blue solution (Thermo Fisher Scientific) prepared in DPBS with a TC10™ automated cell counter (Bio-Rad Laboratories). The required volume of cell suspension was then adjusted to a final density of 125,000 cells/mL. A total of 25,000 HEK293-D2R cells were seeded into each well of a black, clear, flat-bottom 96-well ibiTreat microscopy plate (Ibidi). The cells were allowed to adhere overnight (17–26 h) in an incubator at 37.0°C under a 5% CO₂ atmosphere.

### Fluorescence microscopy assay

A Cytation™ 5 multimode plate reader and microscope (BioTek Instruments) with a 20× LUCPLFLN objective (Olympus) was used for all imaging experiments. The cells were imaged in a bright-field (BF) channel using parameters: LED intensity = 7, camera gain = 24, integration time = 40 ms. Three different fluorescence channels were used: 531 (40) nm excitation and 593 (40) nm emission filters for RFP (LED intensity = 6, camera gain = 24, integration time = 200 ms), 586 (40) nm excitation and 647 (40) nm emission filter for Texas Red (LED intensity = 3, integration time = 200 ms, camera gain = 24), 628 (40) nm excitation and 705 (40) nm emission filter for Cy5 (LED intensity = 3, integration time = 300 ms, camera gain = 24). These imaging parameters were kept constant throughout the entire experimental series. Data acquisition was performed using Gen5 Image+ v3.12 software (BioTek Instruments), and all measurements were conducted at 37 °C in a 5% CO₂ atmosphere.

To identify fluorescent ligands suitable for D_2_ receptor assays, an initial experiment was performed using 12 fluorescent ligands designed for dopamine receptors available within the research group: FAUC-346-TAMRA^41^ (University of Tartu), CELT-240, CELT-241, CELT-383, CELT-385, CELT-419 (Celtarys Research), FGH-460-2, FGH-476-1, FGH-476-4 (University of Erlangen-Nuremberg), NAPS-Cy3B, NAPS-Atto610, NAPS-TAMRA (University of Tartu). Information on the fluorescence spectra and fluorophores of the ligands used in this study is provided in **Table 1**. For measuring the excitation and emission spectra of the fluorescent ligands, a Synergy™ NEO multimode microplate reader (BioTek Instruments) was used with a black 384-well microscopy plate with a clear flat bottom (384 Well Microplate, Greiner). Data acquisition was performed using Gen5 v2.09 software (BioTek Instruments). For spectral measurements, fluorescent ligands were diluted in Dulbecco’s phosphate-buffered saline (DPBS; Corning) to obtain a final concentration of 300 nM in each well. The chemical structures of commercially available fluorescent ligands are not publicly available, whereas the structures of all other ligands are presented in **Supplementary Fig. S1**.

**Table 1.** Excitation and emission maxima of the fluorescent ligands, together with information on the fluorescent label. Excitation and emission wavelengths were determined experimentally.

| **Fluorescent ligand:** | **Excitation**<br>**wavelength, nm** | **Emission**<br>**wavelength, nm** | **Fluorophore** |
| --- | --- | --- | --- |
| FAUC-346-TAMRA | 560 | 582 | TAMRA |
| CELT-240 | 598 | 629 | BDT-TR<sup>42</sup> |
| CELT-241 | 654 | 674 | Cy5<sup>42</sup> |
| CELT-383 | 602 | 628 | BDT-TR |
| CELT-385 | 594 | 628 | BDT-TR |
| CELT-419 | 562 | 578 | Cy3B<sup>42</sup> |
| FGH-460-2 | 651 | 676 | Cy5 |
| FGH-476-1 | 656 | 680 | Cy5 |
| FGH-476-4 | 656 | 682 | Cy5 |
| NAPS-Atto610 | 616 | 638 | Atto610 |
| NAPS-Cy3B | 560 | 576 | Cy3B<sup>15</sup> |
| NAPS-TAMRA | 558 | 590 | TAMRA |

For the initial fluorescent ligand binding screen, binding to dopamine D_2_ receptor was monitored over time. HEK293-D2R cells were seeded at 20,000 cells/well, allowed to adhere overnight, and then treated with different fluorescent ligands at a final concentration of 10 nM, prepared in pre-warmed phenol red-free DMEM (Carl Roth). To assess nonspecific binding and initiate ligand dissociation, spiperone was added to a final concentration of 10 μM. Fluorescent ligand binding to the receptor was monitored for 2 h, followed by dissociation for 6-7 h. The imaging was done in BF and across all three fluorescence channels (RFP, Texas Red, Cy5). Of all the fluorescent ligands tested, those exhibiting the highest signal-to-noise ratios were selected for saturation binding studies, and the most suitable filter block was determined for each ligand.

For saturation binding, HEK293-D2R cells were seeded at 25,000 cells/well and allowed to adhere overnight (37 °C, 5% CO₂). Cells were then incubated with fluorescent ligand serially diluted 2.5-fold (0-10 nM) in pre-warmed phenol red-free DMEM, in the presence or absence of 10 μM spiperone to define nonspecific binding. Binding was monitored for 5 h across three independent experiments.

For competition binding assays, cells seeded at 25,000 cells/well were incubated with fluorescent ligand (CELT-419, FAUC-346-TAMRA, NAPS-Cy3B, or FGH-460-2; 2 nM final concentration) together with unlabeled competitor serially diluted eightfold (haloperidol, 0-1,000 nM; dopamine, 0-1,000 µM) in pre-warmed phenol red-free DMEM. Displacement of the fluorescent ligand was monitored for 5 h across three to four independent experiments.

For dissociation kinetic assays, cells were incubated with 10 nM final concentration of fluorescent ligand (FAUC-346-TAMRA, CELT-419, NAPS-Cy3B, FGH-460-2, CELT-383, CELT-385, CELT-241, or CELT-240) in pre-warmed phenol red-free DMEM, in the presence or absence of 10 μM spiperone to define nonspecific and total binding respectively. Cells were incubated with the ligands for 2 h at 37 °C in a 5% CO₂ atmosphere. Dissociation was initiated by adding 2 µL of spiperone to each well (10 µM final concentration), and the decrease in fluorescence was monitored for 6-7 h. Data are from three independent experiments.

## Computer vision

### Cell and anomaly segmentation models

For the development of the cell segmentation model, a previously constructed dataset^15^ of BF microscopy images of HEK293 cells expressing dopamine D_3_ receptors acquired as Z-stacks was used. The complete dataset formed a data cube with dimensions of 64 × 12 × 904 × 1,224 pixels, corresponding to field of view (FOV) × Z-plane × image height (Y) × image width (X). A total of 64 image stacks were collected from 16 different wells, with consecutive focal planes separated by 4 μm. Each image had dimensions of 904 × 1,224 × 1 pixels, corresponding to a physical area of 292 μm × 395 μm.

Microscopy images were manually annotated using Microsoft Paint. In total, 15 BF images from different FOVs were annotated and subsequently divided into training, validation and test datasets (12, 2, and 1, respectively). During annotation, three principal structures were identified in each image: the cell interior, the cell membrane, and the background adjacent to the cell (**Fig. 1**).

**Figure 1.**
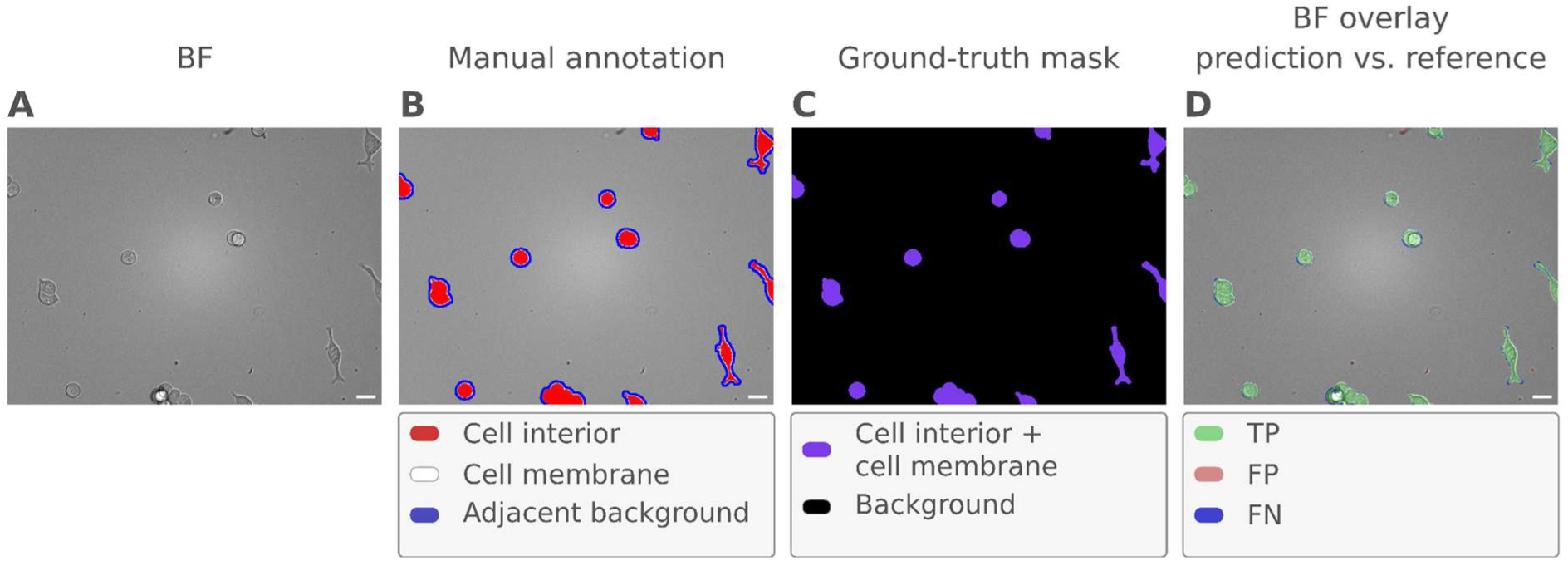
Segmentation quality of the bright-field (BF) cell segmentation model compared with manually annotated reference regions. (A) Original BF microscopy image of HEK293-D3 cells from the test set. (B) Manual annotation of the same field of view, distinguishing the cell interior, cell membrane, and membrane-adjacent background. (C) Binary ground-truth segmentation mask generated from the manual annotation by combining the cell interior and membrane into a single cell class, while the remaining pixels were assigned to the background class. (D) Comparison of the cell segmentation model prediction with the reference ground-truth regions overlaid on the original BF image. True-positive (TP) regions, corresponding to correctly segmented cell pixels, are shown in green; false-positive (FP) regions, corresponding to pixels incorrectly classified as cells, are shown in red; and false-negative (FN) regions, corresponding to reference cell pixels missed by the model, are shown in blue.

Based on the annotated images, binary ground-truth segmentation masks (904 × 1,224 × 1 pixels) were generated, in which pixels corresponding to the cell interior and membrane were combined into a single foreground class while background and membrane-adjacent background pixels were combined into a single background class (**Fig. 1**). For each annotated FOV image, the corresponding 12-plane Z-stack was used to generate overlapping triplets of consecutive focal planes (Z0–Z2, Z1–Z3, …, Z9–Z11), resulting in 10 three-channel input images (904 × 1,224 × 3 pixels) per FOV. Each resulting input image was paired with the ground-truth segmentation mask of the respective FOV, giving 120 training, 20 validation and 10 test input image–mask pairs. Combining multiple focal planes provided complementary morphological information and improved the robustness of cell segmentation to focus variations, enabling cells to be identified even when they were not sharply focused in a single BF image.

Before being combined into three-channel input images, each raw BF focal plane image was individually background-corrected by subtracting a fitted third-degree polynomial surface and adding back its mean. All three-channel input images and ground-truth masks were then padded to 1,248 × 1,248 [H × W] pixels (mean pixel value for images, zeros for masks). To increase model robustness, the 120 training image–mask pairs were augmented with horizontal flips, vertical flips, transpositions, 90° rotations, brightness adjustment and multiplicative intensity noise. Each pair was augmented 25 times with a random combination of these transformations, giving 3,000 training image–mask pairs. The augmented dataset was used to train a U-Net-inspired^43^ convolutional neural network (U-Net-CNN). The network took 1,248 × 1,248 × 3 [H × W × Z (focal plane)] px input images and was trained with binary cross-entropy loss using the Adam optimizer^44^ (learning rate = 2 × 10⁻⁴, β₁ = 0.9, β₂ = 0.999, ε = 10⁻⁸). The model converged within 15 epochs, and its segmentation performance on the test FOV was considered sufficient for the subsequent analysis.

To exclude image regions containing cell debris, fibers or other non-cell objects (hereafter, anomalies), a separate anomaly segmentation model was developed. For this, BF microscopy images of HEK293-D2R cells containing anomalies were taken from the initial fluorescent ligand binding screen. The dataset formed a data cube with dimensions of 191 × 3 × 904 × 1,224 pixels, corresponding to FOV × Z-plane × image height (Y) × image width (X). The 191 FOVs were collected from 48 different wells, with consecutive focal planes separated by 4 μm. Each focal plane image had dimensions of 904 × 1,224 × 1 pixels, corresponding to a physical area of 292 μm × 395 μm.

Anomaly regions were manually annotated to produce binary segmentation masks (904 × 1,224 × 1 pixels) distinguishing anomalies from background. To avoid data leakage, the annotated FOVs were divided into training, validation and test datasets at the well level (131 FOVs from 33 wells, 28 FOVs from 7 wells and 32 FOVs from 8 wells, respectively), so that all FOVs from a given well were assigned to the same dataset. Each raw BF focal plane image was background-corrected as described for the cell segmentation model. The three focal planes of each FOV were then combined into a single three-channel input image (904 × 1,224 × 3 pixels) and paired with the ground-truth mask of the respective FOV.

All three-channel input images and ground-truth masks were then padded to 1,248 × 1,248 [H × W] pixels (mean pixel value for images, zeros for masks). To increase model robustness, the 131 training image–mask pairs were augmented by separately applying horizontal flips, vertical flips, random 90° rotations and transpositions. Each image and its corresponding mask was thus augmented four times, giving 655 training image-mask pairs together with the originals. The augmented dataset was used to train a U-Net-CNN with the same architecture and training settings as the cell segmentation model. The model converged within 25 epochs, and its segmentation performance was considered sufficient for the subsequent analysis. It was then applied to all BF images acquired in this study for automated segmentation of anomalies.

### Cell segmentation and image quantification

For each time point, four regions of interest (ROIs) were imaged in every well. Images were acquired at three different focal planes in both the BF and fluorescence channels. Since cells are visible in the BF images, the BF channel was used as the input for the machine-learning model employed for cell segmentation. In the fluorescence channel, only the pixel intensities corresponding to regions identified as cells by the cell segmentation model were quantified. Background signal was estimated as the mean fluorescence intensity of pixels classified as background by the model. Furthermore, it can be demonstrated that, upon addition of a competing ligand, the fluorescent ligand no longer binds to receptors located on the cell membrane, resulting in a low fluorescence signal at the cell surface.

Microscopy images were analyzed using the MembraneTools module of the Aparecium software platform. Focal planes and region masks were generated using both a cell segmentation model and an anomaly segmentation model. The anomaly segmentation model identified image regions containing cell debris, fibers, or other non-cell objects that could fluoresce or interfere with accurate cell segmentation. These regions were excluded from further analysis because the fluorescence intensity measured in such areas does not originate from living cells and may therefore artificially increase or decrease the signal. For the cell segmentation model, a probability threshold of 0.5 was applied, and all cell pixels (membrane and cytosol) were included in the analysis. For the anomaly segmentation model, a probability threshold of 0.3 was used.

Using MembraneTools, wells were linked to the corresponding concentrations of added compounds as previously described^45^. For each well at each time point, a weighted average of the foreground (cell) and background fluorescence intensities across the four ROIs was calculated, using the corresponding cell or background pixel counts as weights. All experimental data were stored in MIDAS format to ensure unambiguous and experimenter-independent interpretation of the results. Following image analysis, the final background-corrected fluorescence intensity was calculated as:

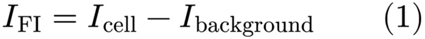

where *I_FI_* is the background-corrected fluorescence intensity in a well, *I_cell_* is the fluorescence intensity associated with the cells, and *I_background_* is the background fluorescence intensity. The resulting data were exported and analyzed using GraphPad Prism 8.0.2 (GraphPad Software).

### Statistical analysis

The suitability of fluorescent ligands for the D_2L_ receptor was evaluated using the Z′-factor, calculated as:

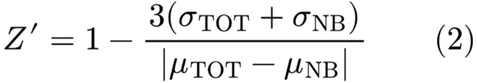

where *σ_TOT_* represents the standard deviation of total binding signal and *σ_NB_* the standard deviation of nonspecific binding, which in this study was assumed to be negligible due to *σ* scaling heteroscedastically with the signal and non-specific binding signal being essentially close to zero in all cases. *μ_TOT_* denotes the mean total binding signal, while *μ_NB_* denotes the mean nonspecific binding signal.

Saturation binding data were analyzed using the built-in “One site – Total and nonspecific binding” model in GraphPad Prism. For CELT-419, the B_max_ parameter was fixed at 25,000 based on two association experiments that lasted for longer (4 h) and where plateau had clearly formed. Competition binding data were analyzed using the built-in “One site – Fit logIC_50_” model. Based on the competition binding results, the equilibrium inhibition constant of the competitor (Kᵢ) was calculated using the Cheng–Prusoff equation:

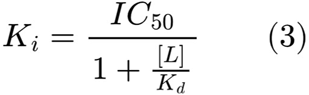

where *IC_50_* is the fitted half-maximal inhibitory concentration, *[L]* is the equilibrium concentration of the fluorescent ligand and *K_d_* is its dissociation constant.

## Results and Discussion

### Selection of fluorescent ligands binding to the D2 receptor

Selecting a suitable fluorescent ligand for binding assays involves navigating a complex trade-off between different properties beyond *K_d_* while the relative importance of each depends on the intended application. In practice, the choice is often limited to a small number of candidates, if any are available at all. Here, we assembled a panel of 12 fluorescent ligands previously characterized at the D_3_ receptor and considered as candidates for repurposing as the D_2_ receptor fluorescent ligands. Working with a relatively large panel allowed us to establish and apply a more rigorous selection framework. Several properties are relevant depending on the application, including signal-to-noise ratio, molecular brightness and quantum yield, excitation and emission spectra, binding affinity, association and dissociation kinetics, extent of nonspecific interactions, photostability, and commercial availability. We, therefore, aimed to select the most suitable fluorescent ligands for the D_2_ receptor by evaluating these properties in the context of typical assay requirements.

Because most of these candidate ligands lack characterization at the D_2_ receptor, we next assessed their binding using an automated fluorescence microscopy-based assay. Binding data for several of the fluorescent ligands tested are available in the literature for the D_3_ receptor, whereas for the D_2_ receptor such data are either lacking or indicate comparatively low affinities (e.g., in radioligand binding assays, CELT-419 has reported K_i_ of approximately 151.4 nM). However, our preliminary experiments indicated that the affinity could be higher than reported in the literature. We therefore screened all available fluorescent ligands for binding to the D_2_ receptor.

We first set out to qualitatively describe interactions between the fluorescent ligand panel and D_2_ receptor expressing HEK293 cells (HEK293-D_2_R), as it allowed us to identify probes for which binding was clearly absent under the tested experimental conditions (**Fig. 2, Supplementary Fig. S2**). In total, 12 different fluorescent ligands were evaluated across three fluorescence channels (RFP, Texas Red, and Cy5) to assess their qualitative binding to the D_2_ receptor **(Supplementary Fig. S3)**. We then confirmed these observations and selected suitable ligands based on their Z-prime metric **(Supplementary Table S1)**.

**Figure 2.**
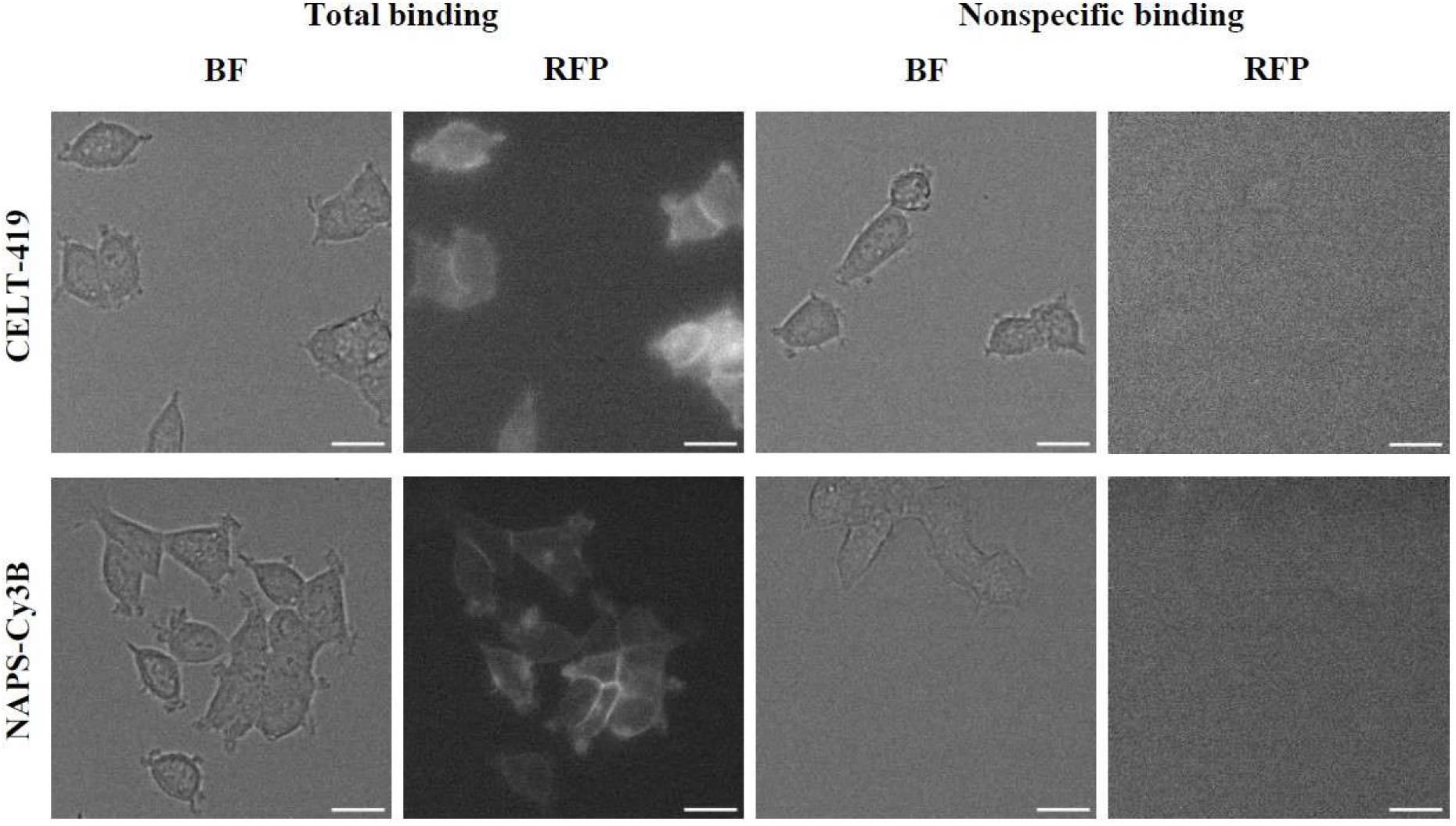
Bright-field (BF) and corresponding fluorescence microscopy images (RFP) of CELT-419 (upper row) or NAPS-Cy3B (lower panel) binding to HEK293-D2R cells. HEK293-D2R cells were incubated with 10 nM fluorescent ligand solution for 2 h. Nonspecific binding was determined in the presence of 10 μM spiperone. The scale bar corresponds to 20 μm. For better visualization, the displayed image’s intensity range was set according to the minimum and maximum pixel values, using Fiji software’s default visualization settings.

For quantification, we used a modified version of a deep-learning based image-analysis pipeline developed previously for this type of assays. Four ligands showed no measurable separation between the total binding signal and image background, nor between total and nonspecific binding conditions. Together with previous D_3_ receptor binding data, this suggests that these four ligands are selective for the D_3_ receptor over D_2_ receptor, making them useful candidates for assays in which D_3_ selectivity is required. The other eight ligands (FAUC-346-TAMRA, CELT-240, CELT-241, CELT-383, CELT-385, CELT-419, FGH-460-2, and NAPS-Cy3B) showed a clear difference between total and nonspecific binding conditions and were, therefore, selected for further study. We then performed saturation binding assays in live cells to determine binding affinities and kinetic parameters, and to establish suitable optical properties and imaging conditions for each ligand.

### Saturation binding assays

The kinetic assay setup enabled monitoring saturation binding over several hours (**Fig. 3**) allowing determination of kinetic behaviour as well as identification of the incubation period required for the signal to reach a plateau (**Fig. 3** and **Fig. 4**). For the eight tested fluorescent ligands, fluorescence intensity increased as ligand concentration increased, indicating growing occupancy of the receptor pool. Most radioligand binding assays are measured after 1 h incubation^45,46^, meaning that the affinities determined in those experiments may be underestimated. Indeed, examining the ligands’ association kinetics over time revealed that a 1 h incubation was clearly not sufficient to reach equilibrium. For some ligands like CELT-383 and CELT-385, no plateau was observed at all during the measurement window at any concentration (**Fig. 3**), suggesting that longer measurement times or higher ligand concentrations may be required for the ligand-receptor complex to reach equilibrium. Surprisingly, two different kinetic behaviours were observed. In most cases, the data exhibited a conventional exponential decay profile indicating the law of mass action type process. In contrast, compounds like CELT-383, CELT-385 and CELT-419, displayed a pronounced lag phase suggesting either a more complex mechanism or limited compound solubility. The latter is unlikely, however, as insoluble conglomerates are usually visible as bright particles in microscopy images, which were not observed. Furthermore, the duration of the lag phase seems to be concentration dependent, introducing additional complexity to the kinetic analysis. Importantly, as the observed association rate constant k_obs_ depends on fluorescent ligand concentration for some ligands, while others display an apparent concentration-dependent lag phase, no single time to equilibrium applies across the panel. Instead, this parameter should be interpolated or extrapolated from the available data for each ligand. Unlike the association phase, dissociation curves display the simplest single-phase exponential decay for all 8 fluorescent ligands (**Fig. 3 I, J**) allowing calculation of k_off_ values for all fluorescent ligands. The k_off_ values were (1.9 ± 0.3) × 10^-2^ min^-1^ for FAUC-346-TAMRA, (2.3 ± 1.1) × 10^-2^ min^-1^ for CELT-240, (3.2 ± 1.8) × 10^-2^ min^-1^ for CELT-241, (2.8 ± 1.4) × 10^-2^ min^-1^ for CELT-383, (0.9 ± 0.5) × 10^-2^ min^-1^ for CELT-385, (1.3 ± 0.4) × 10^-2^ min^-1^ for CELT-419, (3.6 ± 0.5) × 10^-2^ min^-1^ for FGH-460-2 and (1.25 ± 0.09) × 10^-2^ min^-1^ for NAPS-Cy3B.

**Figure 3.**
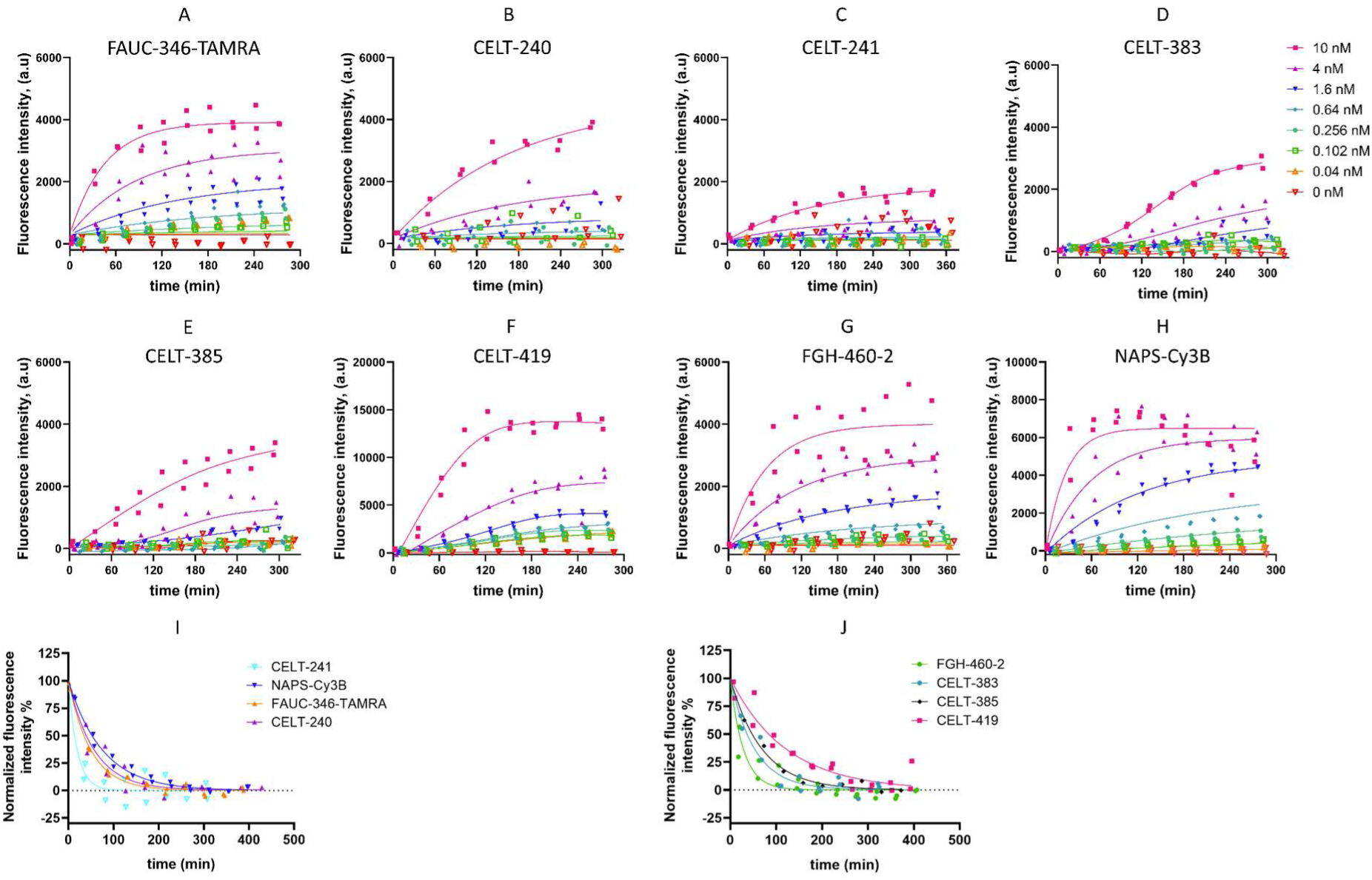
Association and dissociation kinetics of fluorescent ligand binding in HEK293-D2R cells. A varying concentration of 8 fluorescent ligands (**A** - FAUC-346-TAMRA; **B** - CELT-240; **C** - CELT-241; **D** - CELT-383; **E** - CELT-385; **F** - CELT-419; **G** - FGH-460-2; **H** - NAPS-Cy3B) was added to HEK293-D2R cells and measured at approximately 30 minute intervals during association phase. Measurements were continued for approximately 5 h. Two different fits were added to data to guide the eye: One phase association with globally shared parameters (**A-C, G-H**) when no lag phase was observed or smoothing spline (**D-F**) where a lag phase was seen for at least one fluorescent ligand concentration. During the dissociation experiments (**I**, **J**) 10 nM concentration of each fluorescence ligand was added to four parallel wells incubated for 2 h. Thereafter, dissociation was initiated in two parallel wells using 10 µM spiperone while keeping two parallel wells as total binding stability controls (not shown). Dissociation was followed for approximately 7 h. In dissociation (I and J), data is normalized so the exponential decay has Y_0_ of 100% and plateau of 0%, 100% value was fixed to measured fluorescence intensity value of the total binding control values at the beginning of dissociation each replicate was normalized separately. Dissociation data was fitted with a single phase exponential decay function. As each replicate was measured at a slightly different timepoint due to imaging speed, replicate wells have small temporal offsets. Each condition was measured in duplicates in 3 independent experiments.

**Figure 4.**
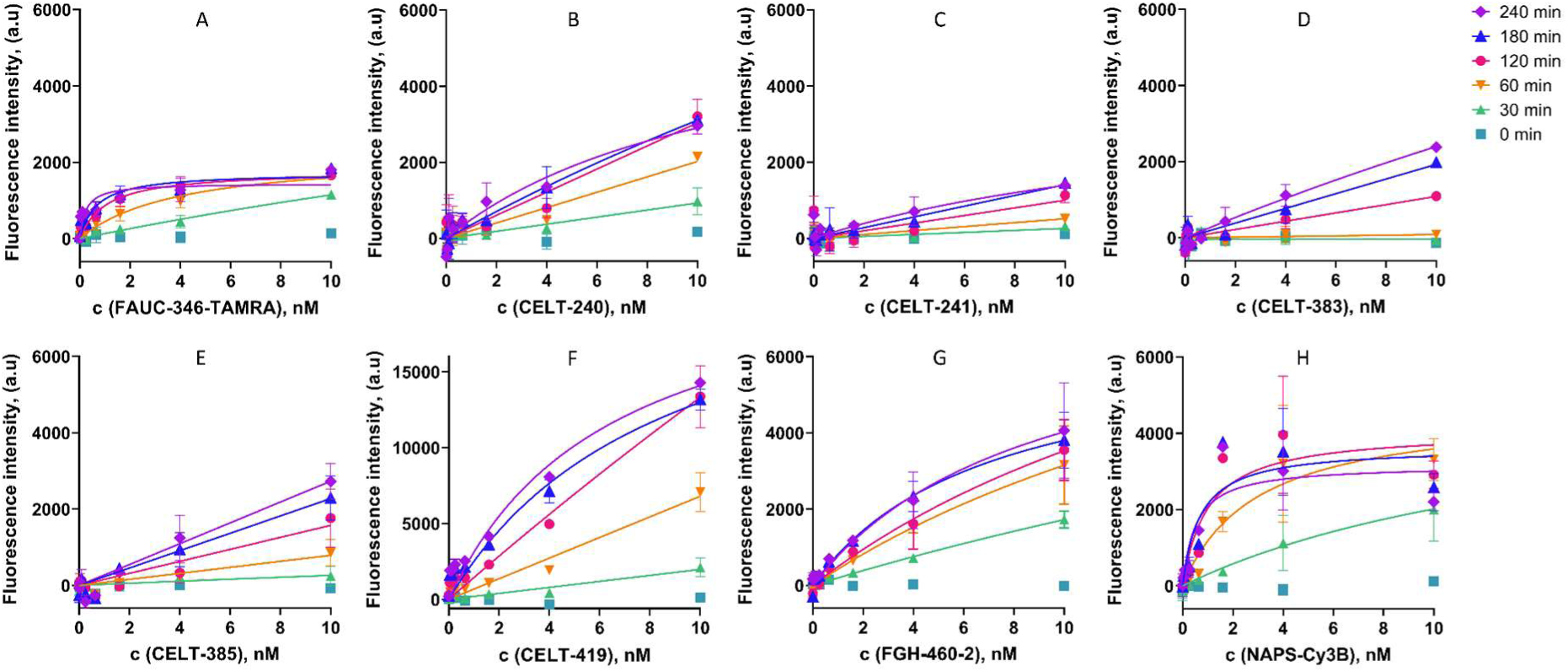
Specific saturation binding curves for the fluorescent ligands FAUC-346-TAMRA (A), CELT-240 (B), CELT-241 (C), CELT-383 (D), CELT-385 (E), CELT-419 (F). FGH-460-2 (G) or NAPS-Cy3B (H) measured over time (0 ‒ 240 min). Saturation binding experiments were performed with live HEK293-D2R cells using 0-10 nM fluorescent ligand concentrations. Nonspecific binding was determined in the presence of 10 μM spiperone. Data points show mean ± SD of one representative out of n = 3 experiments, with each condition carried out in duplicate. Note that the y-axis of CELT-419 (F) is different to better visualize the binding curves.

These experiments allow us to estimate K_d_ values for the panel of ligands. For three fluorescent ligands, the results showed a classic hyperbolic specific binding curve (**Fig. 5A, 5G and 5H**) yielding K_d_ ± SEM values of 2.6 ± 1.2 nM for FAUC-346-TAMRA, 6.1 ± 1.3 nM for FGH-460-2, and 2.0 ± 1.1 nM for NAPS-Cy3B after 2 h incubation.

**Figure 5.**
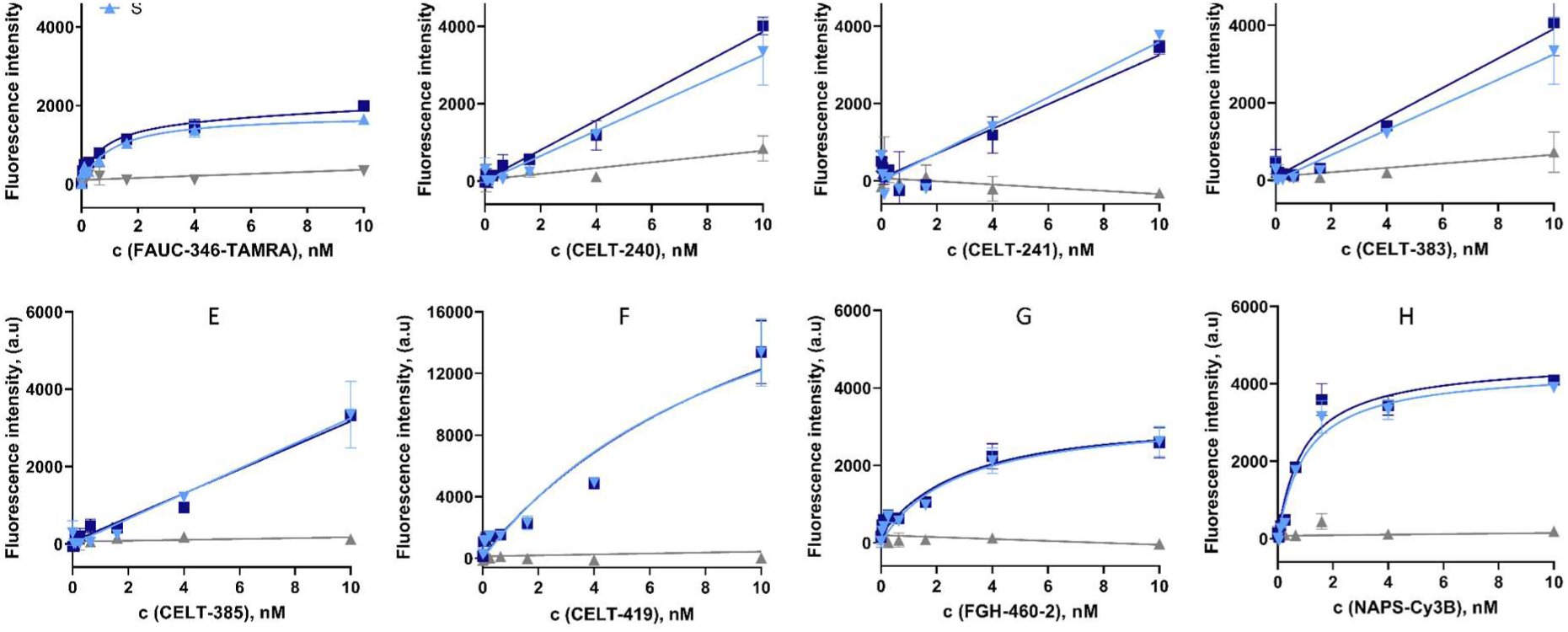
Saturation binding of fluorescent ligands in HEK293-D2R cells. TOT denotes total binding, NB nonspecific binding, and S specific binding of fluorescent ligands FAUC-346-TAMRA (A), CELT-240 (B), CELT-241 (C), CELT-383 (D), CELT-385 (E), CELT-419 (F). FGH-460-2 (G) or NAPS-Cy3B (H) measured after 2 h of incubation. Wells were supplemented with fluorescent ligand solution at varying concentrations (0 ‒ 10 nM using 2.5-fold dilutions). Spiperone was added to the NB wells at a final concentration of 10 μM. Note that the y-axis of CELT-419 (F) is different to better visualize the binding curves. Measurements were performed in duplicate (n = 3) and error bars shown in the figures represent SD.

For the remaining fluorescent ligands, binding curves did not display the expected hyperbolic saturation by the end of the incubation period (**Fig. 5B-F**). Instead, the data followed an approximately linear trend across the tested concentration range. For some ligands, the signal continued to increase even at the final timepoint, whereas for others it appeared to plateau. The effect is explained by the concentration dependent lag phase described earlier. Lower concentrations take longer to equilibrate beyond the measurement window and distort the hyperbolic curve into a near-linear one. In any case, such unsaturated binding curves do not permit K_d_ estimation from concentration-signal curves without prior knowledge of FI_max_, which is itself technically complicated to obtain, owing to the high ligand concentrations required, high background signal, degradation of signal-to-noise ratio, and potential nonspecific uptake. Alternatively, a more complex mechanistic modeling might allow calculating K_d_. For CELT-419, we were nonetheless able to calculate a K_d_ ± SEM value using data from the 3 h timepoint: 17 ± 9 nM. For the remaining ligands, we can only conclude that K_d_ exceeds the half-maximum concentration tested, i.e., K_d_ > 4 nM, although the true affinity could be substantially lower.

### Competition binding assays

After saturation binding assays, we applied the four selected fluorescent ligands with determined affinities (FAUC-346-TAMRA, NAPS-Cy3B, CELT-419, and FGH-460-2) in competition binding assays to determine affinities of unlabeled orthosteric ligands. We selected the endogenous agonist dopamine and the antagonist haloperidol as competing ligands (**Fig. 6-8**). As expected, cellular fluorescence intensity decreased with increasing haloperidol (**Fig. 6**) and dopamine concentrations: higher competitor concentrations led to greater occupancy of the D_2_ receptor orthosteric site by the unlabeled ligand, precluding fluorescent ligand binding and producing a corresponding drop in the measured fluorescence signal. This concentration-dependent signal was readily quantified as described above for the saturation binding assays.

**Figure 6.**
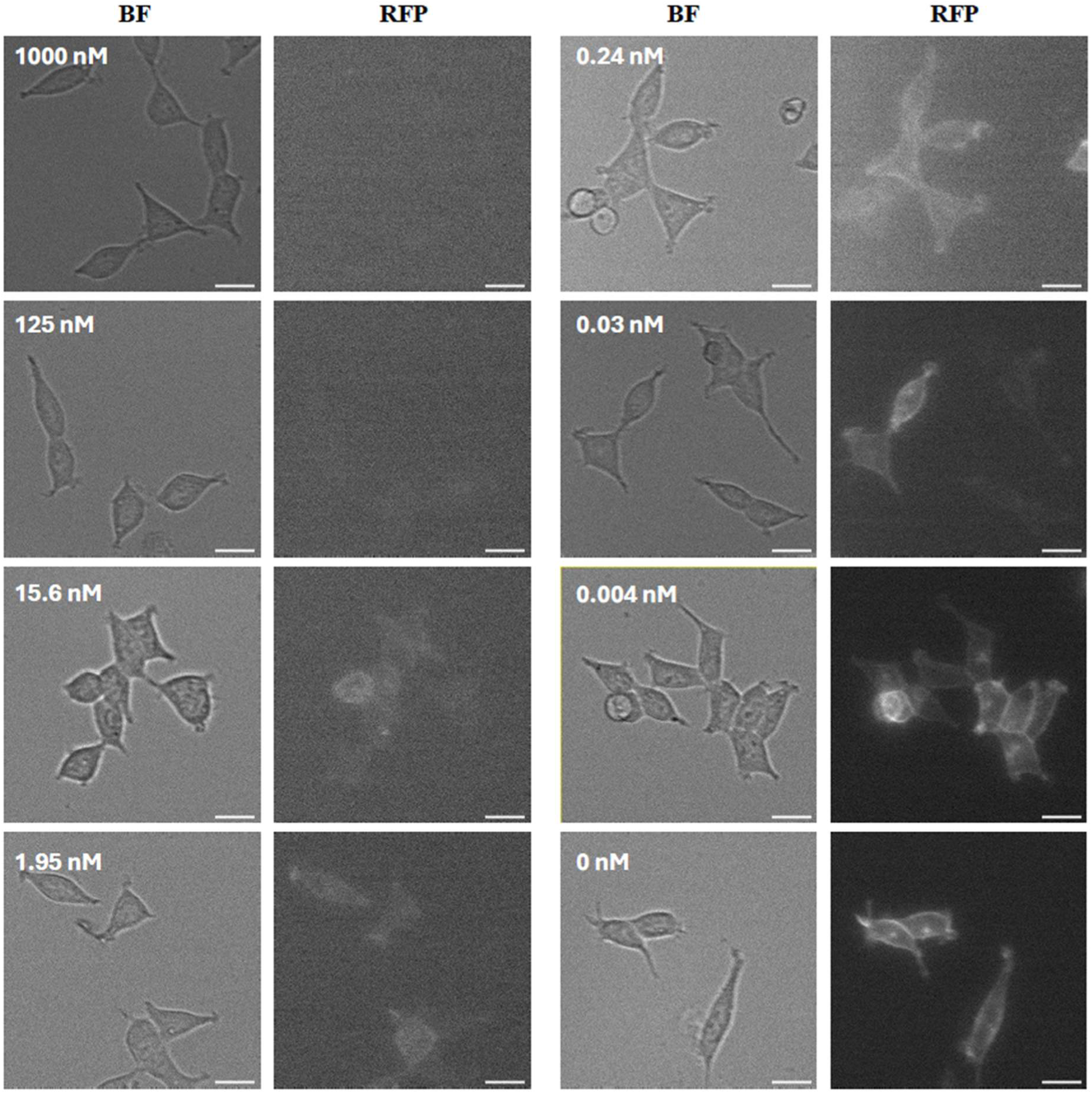
Fluorescence microscopy (RFP) and bright-field (BF) images showing binding of the 2 nM fluorescent ligand NAPS-Cy3B to the D_2_ receptor in the presence of varying concentrations of competing ligand haloperidol after 2 h incubation. Figure shows microscopy images of a representative experiment performed in duplicates (n = 4). The scale bar corresponds to 20 μm. For better visualization, the displayed image’s intensity range was set according to the minimum and maximum pixel values, using Fiji software’s default visualization settings.

To identify equilibrium conditions appropriate for affinity determination, we first characterized the binding kinetics. For all ligands tested, a sigmoidal dose-response relationship developed progressively over time (**Fig. 7**). When testing dopamine, equilibrium was reached faster with FAUC-346-TAMRA (approximately 1 h) than with NAPS-Cy3B, CELT-419 and FGH-460-2 (at least 2 h). With haloperidol, the competition binding kinetics were more similar across probes, requiring at least 2 h of incubation for the change in fluorescence intensity to slow. We then looked at the progression of pIC_50_ values in time to determine if this signal stabilizes faster than the binding process itself and if pre-equilibrium K_i, app_ values serve as reliable proxy for true equilibrium K_i_ values (**Fig. 8C-D**). pIC_50_ values stabilized after approximately 1 h, with no significant further change thereafter for any of the tested probe-competitor pairs. This indicates that although strict equilibrium may not always have been reached within the measurement window, pIC_50_ stabilizes considerably earlier than full equilibrium binding is achieved, such that any resulting bias in the derived affinity estimates is expected to be minor. On this basis, we considered the pIC_50_ (and derived K_i_) values obtained under our standard assay conditions suitable for affinity comparisons between fluorescent probes. Altogether, these results indicate that the time required to reach equilibrium is not fixed, but depends on the intrinsic binding kinetics of both the fluorescent probe and the competing ligand, meaning equilibration time must be empirically determined for each probe-ligand combination rather than assumed from a single reference condition.

**Figure 7.**
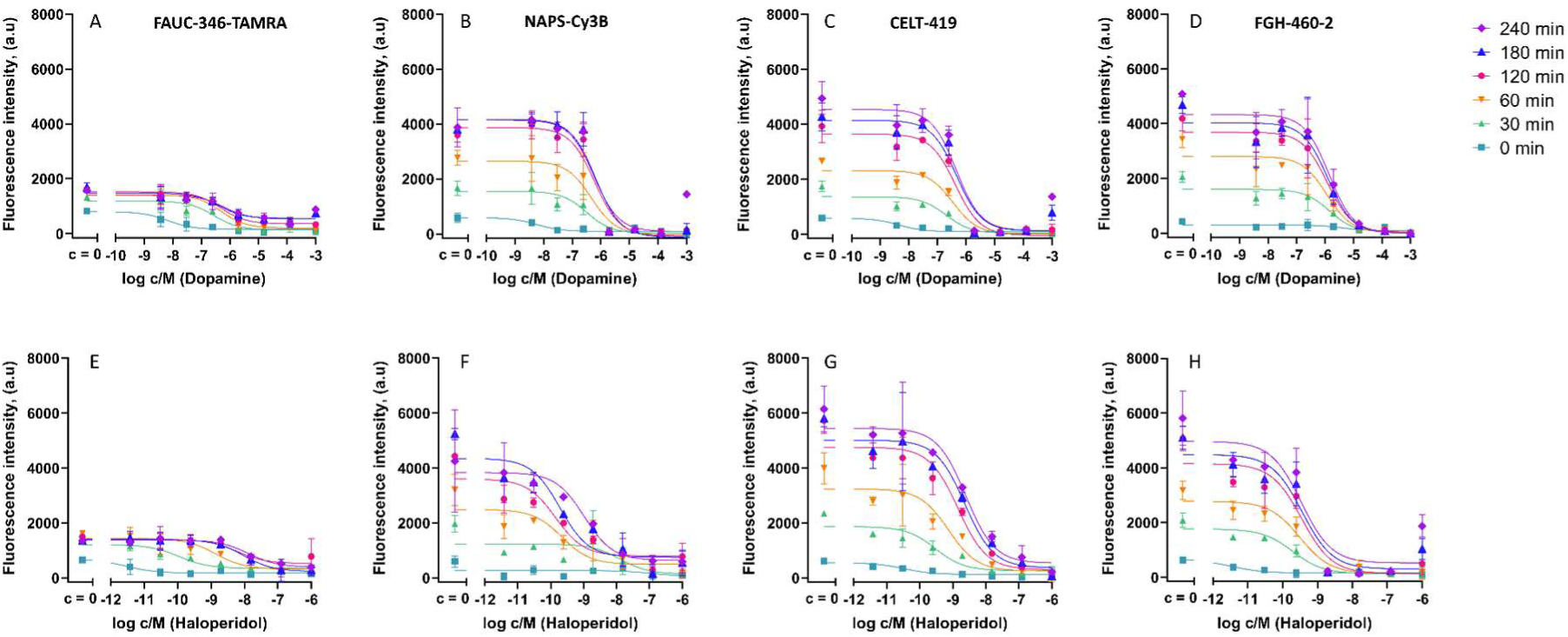
Binding of the fluorescent ligands FAUC-346-TAMRA (panels A and E), NAPS-Cy3B (panels B and F), CELT-419 (panels C and G), and FGH-460-2 (panels D and H) to the dopamine D_2_ receptor in the presence of the competing ligand dopamine (upper row) or haloperidol (lower row). The figures show fluorescence intensity as a function of the added unlabeled ligand concentration at different time points (0 - 240 min). Measurements were performed in duplicate (n = 3-4) and error bars shown in the figures represent SD.

**Figure 8.**
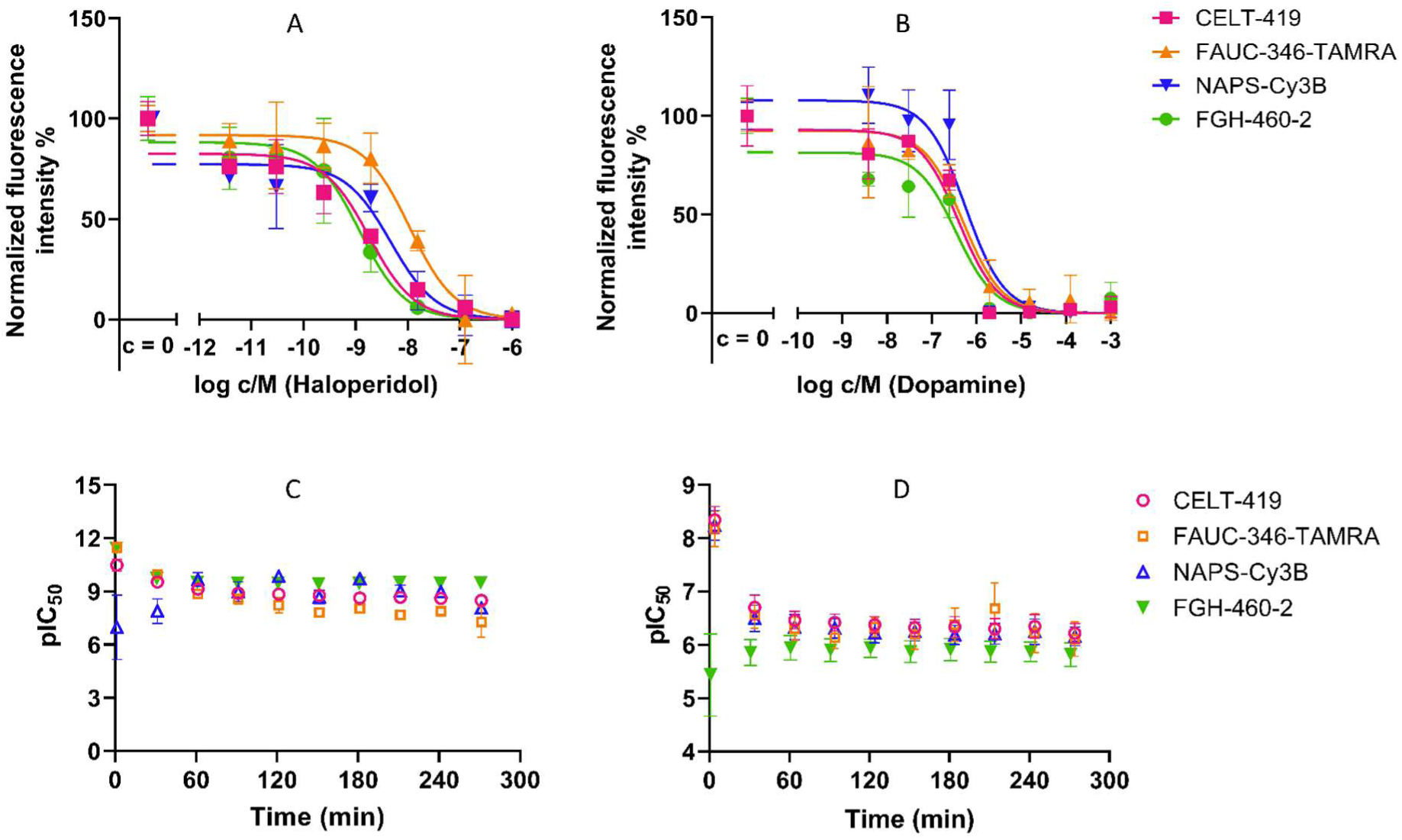
Competition binding of fluorescent ligands with dopamine D_2_ receptor. In the upper row - binding of the 2 nM fluorescent ligands to the D_2_ receptor in the presence of the competing ligand haloperidol (A) or dopamine (B) over 2 h. The x-axis shows the logarithm of the molar concentration. The data were normalized within each dataset: 0% corresponds to the minimum mean signal and 100% to the average FI of the c_competitor_ = 0 experimental points for this particular competitor in the corresponding experiment. For each ligand the experiment was repeated for 3 or 4 times and error bars represent SD. The lower row presents a representative experiment of corresponding pIC_50_ change over time of haloperidol (C) or dopamine (D) determined with different fluorescent ligands. Measurements were performed in duplicate and error bars represent SD. Each experiment was independently repeated (n=3).

With this validated approach in hand, we compared competition binding curves across the four fluorescent ligands (**Fig. 8**). We obtained highly similar competition binding curves across probes when using dopamine as the competing ligand (**Fig. 8B, D**) yielding correspondingly similar pIC_50_ values (**Table 2**). This consistency suggests that, at least for this lower-affinity endogenous agonist, the fluorescent label does not markedly perturb the ligand-receptor interaction being probed. For haloperidol (**Fig. 8A, C**), we observed somewhat greater variability in pIC_50_ values across the fluorescent ligands, though the differences remained modest compared to expected assay variability. This slight probe-dependent variation may reflect subtle differences in the binding kinetics or binding mode of the individual fluorescent probes (see kinetic analysis above), which become marginally more apparent against a high-affinity competitor such as haloperidol than against dopamine.

**Table 2.** pIC_50_ and K_i, app_ values of the competing ligands determined from competition binding assays. pIC_50_ and pK_i, app_ values are given as arithmetic means ± SEM (n = 3-4).

|  | Dopamine |  |  | Haloperidol |  |  |
| --- | --- | --- | --- | --- | --- | --- |
| | $pIC_{50} \pm$<br>SEM | $pK_{i, app} \pm$<br>SEM | $K_{i, app}, nM$ | $pIC_{50} \pm$<br>SEM | $pK_{i, app} \pm$<br>SEM | $K_{i, app}, nM$ |
| **FAUC-346-TAMRA** | $6.20 \pm 0.11$ | $6.40 \pm 0.11$ | 320 | $7.8 \pm 0.4$ | $8.1 \pm 0.4$ | 8.6 |
| **CELT-419** | $6.0 \pm 0.3$ | $6.0 \pm 0.3$ | 980 | $8.65 \pm 0.05$ | $8.70 \pm 0.05$ | 2.02 |
| **FGH-460-2** | $6.2 \pm 0.3$ | $6.3 \pm 0.3$ | 490 | $9.14 \pm 0.17$ | $9.26 \pm 0.17$ | 0.55 |
| **NAPS-Cy3B** | $6.3 \pm 0.06$ | $6.6 \pm 0.06$ | 230 | $9.0 \pm 0.5$ | $9.4 \pm 0.5$ | 0.44 |

K_i, app_ values for dopamine obtained across the different competition binding assays were consistent with one another (**Table 2**) and agreed well with literature estimates, which report a K_i_ range of 63-20,000 nM for dopamine at the D_2_ receptor^47–49^. K_i, app_ values for haloperidol also converged well across the fluorescent probes, with a maximum difference of only 20-fold between ligands, consistent with the literature range of 1.6-40 nM^48,50–52^. The exceptions were FGH-460-2 and NAPS-Cy3B, for which the haloperidol K_i, app_ value fell below the reported range. The generally high affinity values we obtained for haloperidol are consistent with its known high-affinity antagonism at the D_2_ receptor.

Together, these results indicate that the chemical structure of the fluorescent ligand can influence the affinity measured for an unlabeled competitor. The discrepancy in haloperidol K_i, app_ values between FAUC-346-TAMRA and FGH-460-2, in particular, suggests that different fluorescent probes may perturb competitor binding to differing extents, potentially reflecting subtle differences in binding mode or allosteric interplay at the receptor. Clarifying the scope of this effect will require testing a broader panel of unlabeled ligands.

In the developed assay, all eight selected ligands showed clear time-dependent specific binding behaviour. However, the current assay design benefits substantially from statistical averaging over many cells and FOVs for noise reduction and not all of the ligands may therefore be equally suitable for more demanding, high-resolution microscopy applications.

A clear benefit of carrying out such repurposing studies is that even when a fluorescent ligand shows relatively poor affinity to the new target receptor, this outcome itself provides further evidence of the ligand’s specificity for its original target. This is a significant consideration, as most fluorescent probes are designed and validated against a single target receptor, while related receptors are often not thoroughly tested for potential cross-reactivity.

The results of this repurposing approach are especially interesting for the specific case of dopamine D_2_ receptor. Dopamine is a major neurotransmitter in the central nervous system, and through the D_2_ receptor, regulates biological processes such as motor control, reward and motivation, cognition, learning, endocrine signaling, and mood-related processes^1,2,11^. Despite the central pharmacological importance of the D_2_ receptor, relatively few fluorescent probes are available^53^, and substantially fewer have been validated than for the closely related D_3_ subtype. In recent years, fluorescent ligands have been developed for both D_1_-like^54^, as well as for D_2_-like receptors^15,16,55,56^, yet many have not been systematically evaluated at D_2_ receptors. Fluorescent dopamine receptor ligands have enabled diverse applications, including receptor localization in living cells, studies of receptor internalization and trafficking, characterization of ligand binding kinetics, and resonance energy transfer-based binding assays^12,57,58^. Such approaches have been successfully applied to D_1_, D_3_, and D_4_ receptors facilitating investigations of receptor pharmacology in real time, whereas comparable studies at D_2_ have been comparatively scarce because of the limited availability of suitable fluorescent probes.

Beyond its role as a major regulator of normal physiological processes, the D_2_ receptor has also emerged as a compelling drug target. Several studies have identified disease-relevant polymorphisms in the D_2_ receptor, for example in positions Val^96^Ala, Pro^310^Ser, Ser^311^Cys^59^, Ile212Phe, Met374Arg^60^. To date, comparatively little attention has been paid to designing fluorescent probes for genetic variants of receptors. Essentially all of the challenges associated with repurposing a probe from one receptor subtype to another can resurface when adapting an assay from a wild-type receptor to a mutated variant: there is no guarantee that the affinity or kinetic properties of a given fluorescent probe remain suitable for studying the mutant, particularly when the mutation itself alters affinity for common drugs or the endogenous ligand. Having a large panel of fluorescent probes available is, therefore, a significant advantage, as it is less likely that all probes will simultaneously lose affinity when the structural diversity of their pharmacophore structure is high. Conversely, the ability to directly measure the kinetic parameters and affinity of fluorescent probes with well-characterized structures can provide valuable mechanistic and structural insight. For example, it might allow cross-validating hypotheses regarding the structure, structural dynamics and function of the mutated receptor in a more native environment than is accessible through computational studies, crystal structures, or cryo-EM.

## Conclusion

In this study, we demonstrated that fluorescent ligands originally developed and characterized for the dopamine D_3_ receptor can be successfully repurposed as binding probes for the closely related D_2_ receptor, using a microscopy-based live-cell assay. Screening a panel of twelve candidates identified eight ligands with measurable D_2_ receptor binding, four of which yielded reliable K_d_ values from saturation binding, while kinetic analysis revealed that equilibration times vary substantially between probe-competitor combinations and must be determined empirically rather than assumed. Competition binding experiments with dopamine and haloperidol further showed that K_i_ values obtained with different fluorescent probes broadly agree with each other and with literature estimates, while also revealing subtle probe-dependent differences that highlight the importance of validating any given fluorescent ligand for its intended application. Together, these results establish receptor repurposing as a practical strategy for expanding the limited fluorescent toolkit available for the D_2_ receptor, and suggest that this approach, together with the assay platform developed here, may extend readily to other structurally related receptors and their disease-relevant genetic variants.

## Acknowledgements

We would like to thank Dr. Reet Reinart-Okugbeni for the HEK293 cell line stably expressing dopamine D_2_ receptors and Prof. Ago Rinken for valuable scientific advice and helpful discussions. This work was supported by the Estonian Research Council grant (PSG912).

## Author contributions

KO: Writing - Original Draft, Investigation, Formal analysis, Visualization; TL: Formal analysis, Software, Conceptualization, Writing - Original Draft, KH: Software, Supervision, Writing - Original Draft; MI: Investigation, Formal analysis; SON: Investigation, Formal analysis; FG: Investigation, Formal analysis; DT: Investigation, Formal analysis; HRA: Investigation, Formal analysis; MM: Investigation, Conceptualization; EE: Investigation, Conceptualization, Supervision; MH: Methodology, Funding acquisition, Supervision; ES: Methodology, Funding acquisition, Supervision; AA: Conceptualization, Funding acquisition, Supervision, Writing - Original Draft

## References

(1) Beaulieu, J.-M.; Gainetdinov, R. R. The Physiology, Signaling, and Pharmacology of Dopamine Receptors. Pharmacol Rev 2011, 63 (1), 182–217. 10.1124/pr.110.002642

(2) Weiner, D. M.; Brann, M. R. The Distribution of a Dopamine D2 Receptor mRNA in Rat Brain. FEBS Letters 1989, 253 (1–2), 207–213. 10.1016/0014-5793(89)80960-3

(3) Seeman, P. Targeting the Dopamine D2 Receptor in Schizophrenia. Expert Opin Ther Targets 2006, 10 (4), 515–531. 10.1517/14728222.10.4.515

(4) Wang, S.; Che, T.; Levit, A.; Shoichet, B. K.; Wacker, D.; Roth, B. L. Structure of the D2 Dopamine Receptor Bound to the Atypical Antipsychotic Drug Risperidone. Nature 2018, 555 (7695), 269–273. 10.1038/nature25758

(5) de Bartolomeis, A.; Ciccarelli, M.; De Simone, G.; Mazza, B.; Barone, A.; Vellucci, L. Canonical and Non-Canonical Antipsychotics’ Dopamine-Related Mechanisms of Present and Next Generation Molecules: A Systematic Review on Translational Highlights for Treatment Response and Treatment-Resistant Schizophrenia. International Journal of Molecular Sciences 2023, 24 (6), 5945. 10.3390/ijms24065945

(6) Li, P.; Snyder, G. L.; Vanover, K. E. Dopamine Targeting Drugs for the Treatment of Schizophrenia: Past, Present and Future. Curr Top Med Chem 2016, 16 (29), 3385–3403. 10.2174/1568026616666160608084834

(7) Kiss, B.; Laszlovszky, I.; Krámos, B.; Visegrády, A.; Bobok, A.; Lévay, G.; Lendvai, B.; Román, V. Neuronal Dopamine D3 Receptors: Translational Implications for Preclinical Research and CNS Disorders. Biomolecules 2021, 11 (1), 104. 10.3390/biom11010104

(8) Prieto, G. A. Abnormalities of Dopamine D3 Receptor Signaling in the Diseased Brain. J Cent Nerv Syst Dis 2017, 9, 1179573517726335. 10.1177/1179573517726335

(9) Chien, E. Y. T.; Liu, W.; Zhao, Q.; Katritch, V.; Han, G. W.; Hanson, M. A.; Shi, L.; Newman, A. H.; Javitch, J. A.; Cherezov, V.; Stevens, R. C. Structure of the Human Dopamine D3 Receptor in Complex with a D2/D3 Selective Antagonist. Science 2010, 330 (6007), 1091–1095. 10.1126/science.1197410

(10) Lane, J. R.; Chubukov, P.; Liu, W.; Canals, M.; Cherezov, V.; Abagyan, R.; Stevens, R. C.; Katritch, V. Structure-Based Ligand Discovery Targeting Orthosteric and Allosteric Pockets of Dopamine Receptors. Mol Pharmacol 2013, 84 (6), 794–807. 10.1124/mol.113.088054

(11) Missale, C.; Nash, S. R.; Robinson, S. W.; Jaber, M.; Caron, M. G. Dopamine Receptors: From Structure to Function. Physiol Rev 1998, 78 (1), 189–225. 10.1152/physrev.1998.78.1.189

(12) Allikalt, A.; Purkayastha, N.; Flad, K.; Schmidt, M. F.; Tabor, A.; Gmeiner, P.; Hübner, H.; Weikert, D. Fluorescent Ligands for Dopamine D2/D3 Receptors. Sci Rep 2020, 10 (1), 21842. 10.1038/s41598-020-78827-9

(13) Ariano, M. A.; Monsma, F. J.; Barton, A. C.; Kang, H. C.; Haugland, R. P.; Sibley, D. R. Direct Visualization and Cellular Localization of D1 and D2 Dopamine Receptors in Rat Forebrain by Use of Fluorescent Ligands. Proceedings of the National Academy of Sciences 1989, 86 (21), 8570–8574. 10.1073/pnas.86.21.8570

(14) Prokop, S.; Ábrányi-Balogh, P.; Barti, B.; Vámosi, M.; Zöldi, M.; Barna, L.; Urbán, G. M.; Tóth, A. D.; Dudok, B.; Egyed, A.; Deng, H.; Leggio, G. M.; Hunyady, L.; van der Stelt, M.; Keserű, G. M.; Katona, I. PharmacoSTORM Nanoscale Pharmacology Reveals Cariprazine Binding on Islands of Calleja Granule Cells. Nat Commun 2021, 12 (1), 6505. 10.1038/s41467-021-26757-z

(15) Allikalt, A.; Laasfeld, T.; Ilisson, M.; Kopanchuk, S.; Rinken, A. Quantitative Analysis of Fluorescent Ligand Binding to Dopamine D3 Receptors Using Live-Cell Microscopy. The FEBS Journal 2021, 288 (5), 1514–1532. 10.1111/febs.15519

(16) Tahk, M.-J.; Laasfeld, T.; Meriste, E.; Brea, J.; Loza, M. I.; Majellaro, M.; Contino, M.; Sotelo, E.; Rinken, A. Fluorescence Based HTS-Compatible Ligand Binding Assays for Dopamine D3 Receptors in Baculovirus Preparations and Live Cells. Frontiers in Molecular Biosciences 2023, 10. 10.3389/fmolb.2023.1119157

(17) Paton, W. D. M.; Rang, H. P. The Uptake of Atropine and Related Drugs by Intestinal Smooth Muscle of the Guinea-Pig in Relation to Acetylcholine Receptors. Proceedings of the Royal Society of London. Series B. Biological Sciences 1965, 163 (990), 1–44. 10.1098/rspb.1965.0058

(18) Ertl, F. J.; Kopanchuk, S.; Dijon, N. C.; Veikšina, S.; Tahk, M.-J.; Laasfeld, T.; Schettler, F.; Gattor, A. O.; Hübner, H.; Archipowa, N.; Köckenberger, J.; Heinrich, M. R.; Gmeiner, P.; Kutta, R. J.; Holliday, N. D.; Rinken, A.; Keller, M. Dually Labeled Neurotensin NTS1R Ligands for Probing Radiochemical and Fluorescence-Based Binding Assays. J. Med. Chem. 2024, 67 (18), 16664–16691. 10.1021/acs.jmedchem.4c01470

(19) Allikalt, A.; Kopanchuk, S.; Rinken, A. Implementation of Fluorescence Anisotropy-Based Assay for the Characterization of Ligand Binding to Dopamine D1 Receptors. European Journal of Pharmacology 2018, 839, 40–46. 10.1016/j.ejphar.2018.09.008

(20) Rinken, A.; Lavogina, D.; Kopanchuk, S. Assays with Detection of Fluorescence Anisotropy: Challenges and Possibilities for Characterizing Ligand Binding to GPCRs. Trends in Pharmacological Sciences 2018, 39 (2), 187–199. 10.1016/j.tips.2017.10.004

(21) Keller, M.; Mahuroof, S. A.; Hong Yee, V.; Carpenter, J.; Schindler, L.; Littmann, T.; Pegoli, A.; Hübner, H.; Bernhardt, G.; Gmeiner, P.; Holliday, N. D. Fluorescence Labeling of Neurotensin(8–13) via Arginine Residues Gives Molecular Tools with High Receptor Affinity. ACS Med. Chem. Lett. 2019, 11 (1), 16–22. 10.1021/acsmedchemlett.9b00462

(22) Gleixner, J.; Kopanchuk, S.; Grätz, L.; Tahk, M.-J.; Laasfeld, T.; Veikšina, S.; Höring, C.; Gattor, A. O.; Humphrys, L. J.; Müller, C.; Archipowa, N.; Köckenberger, J.; Heinrich, M. R.; Kutta, R. J.; Rinken, A.; Keller, M. Illuminating Neuropeptide Y Y4 Receptor Binding: Fluorescent Cyclic Peptides with Subnanomolar Binding Affinity as Novel Molecular Tools. ACS Pharmacol. Transl. Sci. 2024, 7 (4), 1142–1168. 10.1021/acsptsci.4c00013

(23) Tahk, M.-J.; Torp, J.; Ali, M. A. S.; Fishman, D.; Parts, L.; Grätz, L.; Müller, C.; Keller, M.; Veiksina, S.; Laasfeld, T.; Rinken, A. Live-Cell Microscopy or Fluorescence Anisotropy with Budded Baculoviruses—Which Way to Go with Measuring Ligand Binding to M4 Muscarinic Receptors? 2022. 10.1098/rsob.220019

(24) Lohse, M. J.; Nuber, S.; Hoffmann, C. Fluorescence/Bioluminescence Resonance Energy Transfer Techniques to Study G-Protein-Coupled Receptor Activation and Signaling. Pharmacological Reviews 2012, 64 (2), 299–336. 10.1124/pr.110.004309

(25) Grätz, L.; Laasfeld, T.; Allikalt, A.; Gruber, C. G.; Pegoli, A.; Tahk, M.-J.; Tsernant, M.-L.; Keller, M.; Rinken, A. BRET- and Fluorescence Anisotropy-Based Assays for Real-Time Monitoring of Ligand Binding to M2 Muscarinic Acetylcholine Receptors. Biochimica et Biophysica Acta (BBA) - Molecular Cell Research 2021, 1868 (3), 118930. 10.1016/j.bbamcr.2020.118930

(26) Dale, N. C.; Johnstone, E. K. M.; White, C. W.; Pfleger, K. D. G. NanoBRET: The Bright Future of Proximity-Based Assays. Front. Bioeng. Biotechnol. 2019, 7. 10.3389/fbioe.2019.00056

(27) Stoddart, L. A.; Johnstone, E. K. M.; Wheal, A. J.; Goulding, J.; Robers, M. B.; Machleidt, T.; Wood, K. V.; Hill, S. J.; Pfleger, K. D. G. Application of BRET to Monitor Ligand Binding to GPCRs. Nat Methods 2015, 12 (7), 661–663. 10.1038/nmeth.3398

(28) Waller, A.; Pipkorn, D.; Sutton, K. L.; Linderman, J. J.; Omann, G. M. Validation of Flow Cytometric Competitive Binding Protocols and Characterization of Fluorescently Labeled Ligands. Cytometry 2001, 45 (2), 102–114. 10.1002/1097-0320(20011001)45:2%3C102::AID-CYTO1152%3E3.0.CO;2-Z

(29) Jacobsen, S. E.; Ammendrup-Johnsen, I.; Jansen, A. M.; Gether, U.; Madsen, K. L.; Bräuner-Osborne, H. The GPRC6A Receptor Displays Constitutive Internalization and Sorting to the Slow Recycling Pathway. Journal of Biological Chemistry 2017, 292 (17), 6910–6926. 10.1074/jbc.M116.762385

(30) Beghin, A.; Kechkar, A.; Butler, C.; Levet, F.; Cabillic, M.; Rossier, O.; Giannone, G.; Galland, R.; Choquet, D.; Sibarita, J.-B. Localization-Based Super-Resolution Imaging Meets High-Content Screening. Nat Methods 2017, 14 (12), 1184–1190. 10.1038/nmeth.4486

(31) Barentine, A. E. S.; Lin, Y.; Courvan, E. M.; Kidd, P.; Liu, M.; Balduf, L.; Phan, T.; Rivera-Molina, F.; Grace, M. R.; Marin, Z.; Lessard, M.; Rios Chen, J.; Wang, S.; Neugebauer, K. M.; Bewersdorf, J.; Baddeley, D. An Integrated Platform for High-Throughput Nanoscopy. Nat Biotechnol 2023, 41 (11), 1549–1556. 10.1038/s41587-023-01702-1

32. (32) Dopamine | D2 Receptor | Fluorescent Antagonist | Ligand. Celtarys. https://www.celtarys.com/categoria-producto/dopamine-d2-receptor-family (accessed 2025-04-14)

(33) Nagl, M.; Mönnich, D.; Rosier, N.; Schihada, H.; Sirbu, A.; Konar, N.; Reyes-Resina, I.; Navarro, G.; Franco, R.; Kolb, P.; Annibale, P.; Pockes, S. Fluorescent Tools for the Imaging of Dopamine D2-Like Receptors. ChemBioChem 2024, 25 (2), e202300659. 10.1002/cbic.202300659

(34) Elek, M.; Dubiel, M.; Mayer, L.; Zivkovic, A.; Müller, T. J. J.; Stark, H. BOPPY-Based Novel Fluorescent Dopamine D2 and D3 Receptor Ligands. Bioorganic & Medicinal Chemistry Letters 2022, 59, 128573. 10.1016/j.bmcl.2022.128573

(35) Tabor, A.; Weisenburger, S.; Banerjee, A.; Purkayastha, N.; Kaindl, J. M.; Hübner, H.; Wei, L.; Grömer, T. W.; Kornhuber, J.; Tschammer, N.; Birdsall, N. J. M.; Mashanov, G. I.; Sandoghdar, V.; Gmeiner, P. Visualization and Ligand-Induced Modulation of Dopamine Receptor Dimerization at the Single Molecule Level. Sci Rep 2016, 6, 33233. 10.1038/srep33233

(36) Tomlinson, I. D.; Kovtun, O.; Crescentini, T. M.; Rosenthal, S. J. Biotinylated-Spiperone Ligands for Quantum Dot Labeling of the Dopamine D2 Receptor in Live Cell Cultures. Bioorganic & Medicinal Chemistry Letters 2019, 29 (8), 959–964. 10.1016/j.bmcl.2019.02.024

(37) Fan, L.; Tan, L.; Chen, Z.; Qi, J.; Nie, F.; Luo, Z.; Cheng, J.; Wang, S. Haloperidol Bound D2 Dopamine Receptor Structure Inspired the Discovery of Subtype Selective Ligands. Nat Commun 2020, 11 (1), 1074. 10.1038/s41467-020-14884-y

(38) Hounsou, C.; Baehr, C.; Gasparik, V.; Alili, D.; Belhocine, A.; Rodriguez, T.; Dupuis, E.; Roux, T.; Mann, A.; Heissler, D.; Pin, J.-P.; Durroux, T.; Bonnet, D.; Hibert, M. From the Promiscuous Asenapine to Potent Fluorescent Ligands Acting at a Series of Aminergic G-Protein-Coupled Receptors. J Med Chem 2018, 61 (1), 174–188. 10.1021/acs.jmedchem.7b01220

(39) Killoran, M. P.; Levin, S.; Boursier, M. E.; Zimmerman, K.; Hurst, R.; Hall, M. P.; Machleidt, T.; Kirkland, T. A.; Friedman Ohana, R. An Integrated Approach toward NanoBRET Tracers for Analysis of GPCR Ligand Engagement. Molecules 2021, 26 (10), 2857. 10.3390/molecules26102857

(40) Reinart-Okugbeni, R.; Ausmees, K.; Kriis, K.; Werner, F.; Rinken, A.; Kanger, T. Chemoenzymatic Synthesis and Evaluation of 3-Azabicyclo[3.2.0]Heptane Derivatives as Dopaminergic Ligands. European Journal of Medicinal Chemistry 2012, 55, 255–261. 10.1016/j.ejmech.2012.07.025

(41) Bettinetti, L.; Schlotter, K.; Hübner, H.; Gmeiner, P. Interactive SAR Studies: Rational Discovery of Super-Potent and Highly Selective Dopamine D3 Receptor Antagonists and Partial Agonists. J. Med. Chem. 2002, 45 (21), 4594–4597. 10.1021/jm025558r

42. Dopamine Receptor Family - Celtarys. https://www.celtarys.com/celtarys-products/fluorescent-ligands/gpcr-ligands/dopamine-receptor-family (accessed 2026-09-14)

43. Ronneberger, O.; Fischer, P.; Brox, T. U-Net: Convolutional Networks for Biomedical Image Segmentation. In Medical Image Computing and Computer-Assisted Intervention – MICCAI 2015; Navab, N., Hornegger, J., Wells, W. M., Frangi, A. F., Eds.; Springer International Publishing: Cham, 2015; pp 234–241. 10.1007/978-3-319-24574-4_28

(44) Kingma, D. P.; Ba, J. Adam: A Method for Stochastic Optimization. arXiv January 30, 2017. 10.48550/arXiv.1412.6980

(45) Laasfeld, T.; Tahk, M.-J.; Allikalt, A.; Torp, J.; Grätz, L.; Kopanchuk, S.; Rinken, A. Exploring Muscarinic Acetylcholine Receptor Binding Kinetics with Fluorescence Anisotropy; 2024; pp 113–151. 10.1007/978-1-0716-4015-9_6

(46) Hulme, E. C.; Trevethick, M. A. Ligand Binding Assays at Equilibrium: Validation and Interpretation. British Journal of Pharmacology 2010, 161 (6), 1219–1237. 10.1111/j.1476-5381.2009.00604.x

(47) Burris, K. D.; Pacheco, M. A.; Filtz, T. M.; Kung, M. P.; Kung, H. F.; Molinoff, P. B. Lack of Discrimination by Agonists for D2 and D3 Dopamine Receptors. Neuropsychopharmacology 1995, 12 (4), 335–345. 10.1016/0893-133X(94)00099-L

(48) Freedman, S. B.; Patel, S.; Marwood, R.; Emms, F.; Seabrook, G. R.; Knowles, M. R.; McAllister, G. Expression and Pharmacological Characterization of the Human D3 Dopamine Receptor. J Pharmacol Exp Ther 1994, 268 (1), 417–426.

(49) Sautel, F.; Griffon, N.; Lévesque, D.; Pilon, C.; Schwartz, J. C.; Sokoloff, P. A Functional Test Identifies Dopamine Agonists Selective for D3 versus D2 Receptors. Neuroreport 1995, 6 (2), 329–332. 10.1097/00001756-199501000-00026

(50) MacKenzie, R. G.; VanLeeuwen, D.; Pugsley, T. A.; Shih, Y. H.; Demattos, S.; Tang, L.; Todd, R. D.; O’Malley, K. L. Characterization of the Human Dopamine D3 Receptor Expressed in Transfected Cell Lines. Eur J Pharmacol 1994, 266 (1), 79–85. 10.1016/0922-4106(94)90212-7

(51) Tice, M. A.; Hashemi, T.; Taylor, L. A.; Duffy, R. A.; McQuade, R. D. Characterization of the Binding of SCH 39166 to the Five Cloned Dopamine Receptor Subtypes. Pharmacol Biochem Behav 1994, 49 (3), 567–571. 10.1016/0091-3057(94)90070-1

(52) Sokoloff, P.; Andrieux, M.; Besançon, R.; Pilon, C.; Martres, M. P.; Giros, B.; Schwartz, J. C. Pharmacology of Human Dopamine D3 Receptor Expressed in a Mammalian Cell Line: Comparison with D2 Receptor. Eur J Pharmacol 1992, 225 (4), 331–337. 10.1016/0922-4106(92)90107-7

53. (53) Xu, Z. Development of Fluorescent and Photoactivatable Ligands Targeting the Dopamine D2 Receptor. thesis, Monash University, 2020. 10.26180/12981491.v1

(54) Gómez-Santacana, X.; Boutonnet, M.; Martínez-Juvés, C.; Cimadevila, M.; Catena, J.; Moutin, E.; Roux, T.; Trinquet, E.; Lamarque, L.; Perroy, J.; Prézeau, L.; Zwier, J. M.; Pin, J.-P.; Llebaria, A. A Modular Click Ligand-Directed Approach to Label Endogenous Dopamine D1 Receptors in Live Cells. Commun Chem 2025, 8 (1), 113. 10.1038/s42004-025-01504-3

(55) Leopoldo, M.; Lacivita, E.; Passafiume, E.; Contino, M.; Colabufo, N. A.; Berardi, F.; Perrone, R. 4-[ω-[4-Arylpiperazin-1-Yl]Alkoxy]Phenyl)Imidazo[1,2-a]Pyridine Derivatives: Fluorescent High-Affinity Dopamine D3 Receptor Ligands as Potential Probes for Receptor Visualization. J. Med. Chem. 2007, 50 (20), 5043–5047. 10.1021/jm070721+

(56) Vellaisamy, K.; Li, G.; Ko, C.-N.; Zhong, H.-J.; Fatima, S.; Kwan, H.-Y.; Wong, C.-Y.; Kwong, W.-J.; Tan, W.; Leung, C.-H.; Ma, D.-L. Cell Imaging of Dopamine Receptor Using Agonist Labeling Iridium(Iii) Complex. Chem. Sci. 2018, 9 (5), 1119–1125. 10.1039/c7sc04798c

(57) Tabor, A.; Möller, D.; Hübner, H.; Kornhuber, J.; Gmeiner, P. Visualization of Ligand-Induced Dopamine D2S and D2L Receptor Internalization by TIRF Microscopy. Sci Rep 2017, 7 (1), 10894. 10.1038/s41598-017-11436-1

(58) Dorogan, M.; Namballa, H. K.; Patra, S.; Harding, W. W. Recent Advances in Dopamine Receptor Ligands as Chemical Biology Tools. ACS Omega 2026, 11 (24), 34870–34884. 10.1021/acsomega.6c01474

(59) Cravchik, A.; Sibley, D. R.; Gejman, P. V. Functional Analysis of the Human D2 Dopamine Receptor Missense Variants*. Journal of Biological Chemistry 1996, 271 (42), 26013–26017. 10.1074/jbc.271.42.26013

(60) Rodriguez-Contreras, D.; García-Nafría, J.; Chan, A. E.; Shinde, U.; Neve, K. A. Comparison of the Function of Two Novel Human Dopamine D2 Receptor Variants Identifies a Likely Mechanism for Their Pathogenicity. Biochemical Pharmacology 2024, 228, 116228. 10.1016/j.bcp.2024.116228

