## Supplementary material for "Repurposing dopamine D_3_ receptor fluorescent ligands for D_2_ receptor": Supplementary information_Allikalt.docx

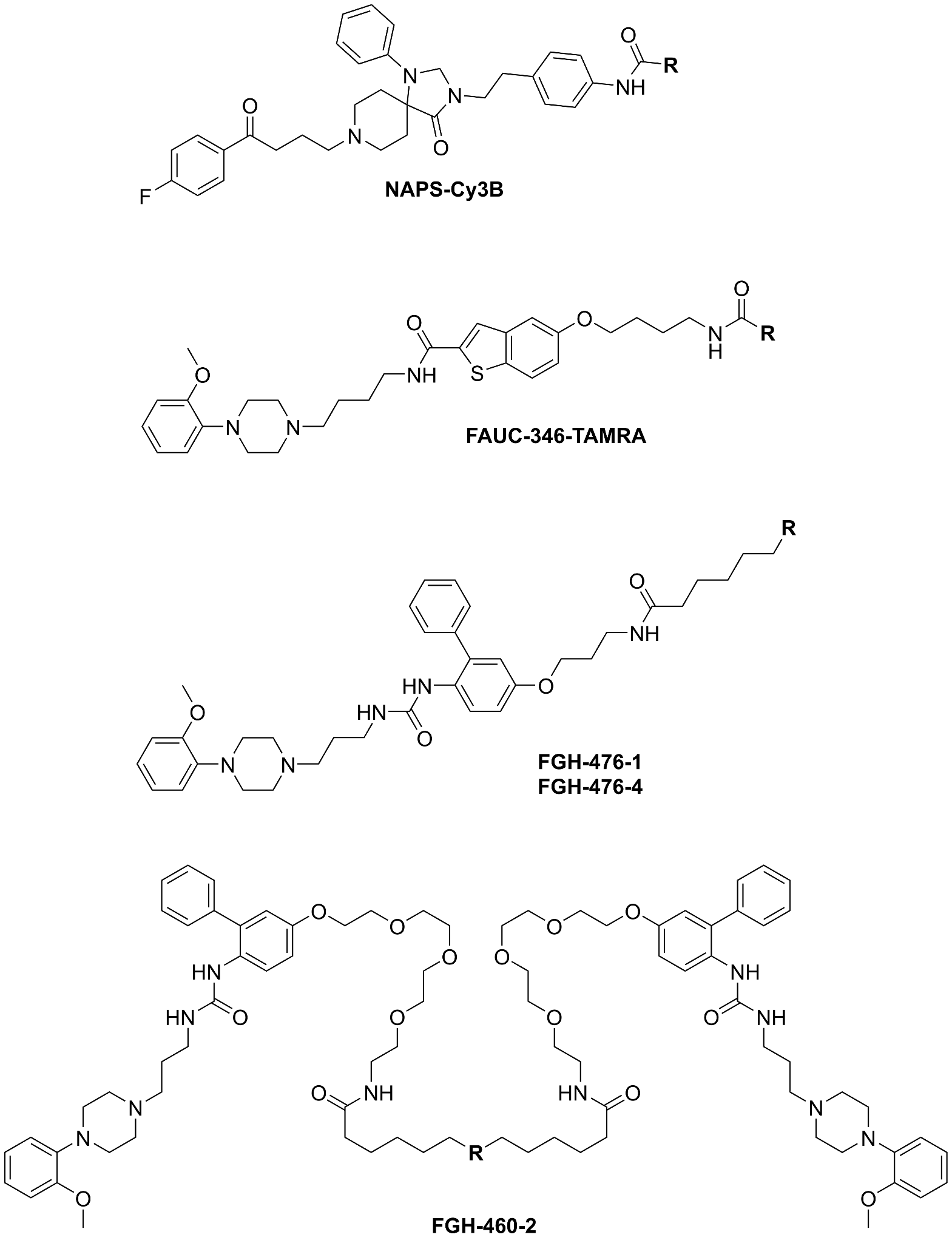

**Supplementary Figure S1**. Structures of fluorescent ligands tested in this manuscript except commercially available fluorescent ligands from Celtarys Research. Fluorophore groups are presented with variable **R**.

A

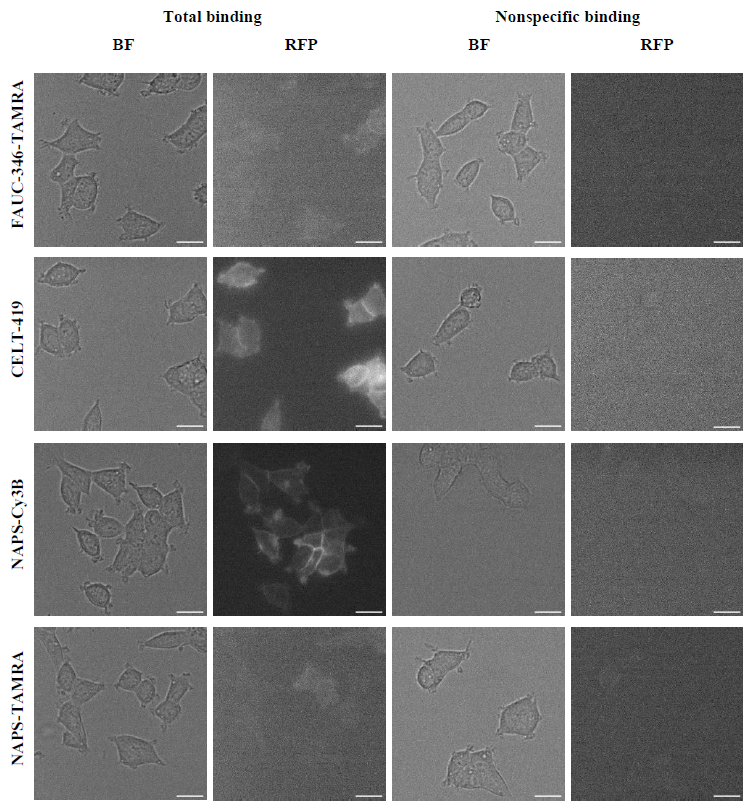

B

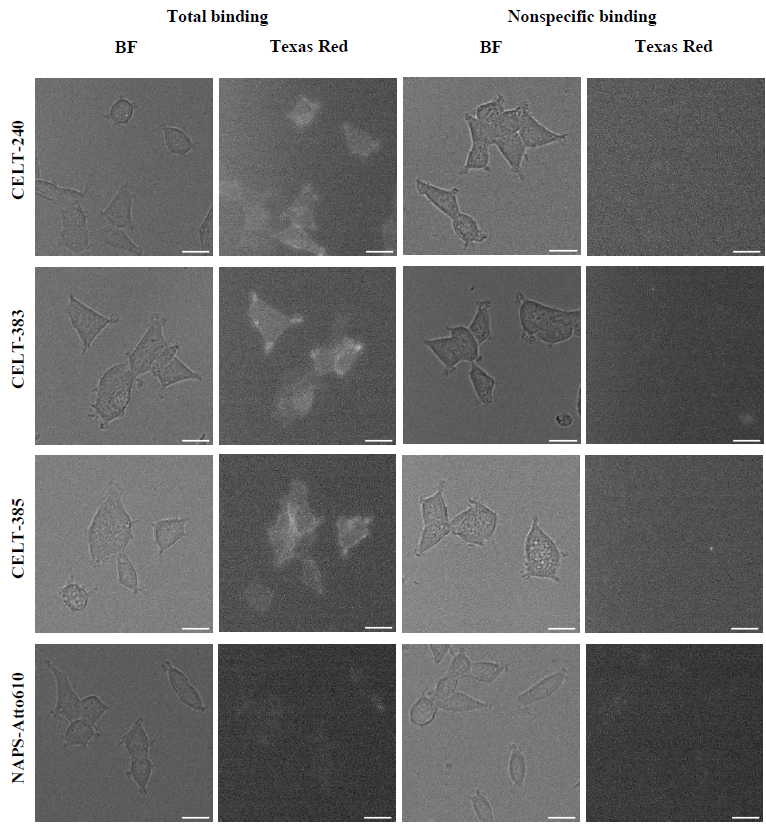

C

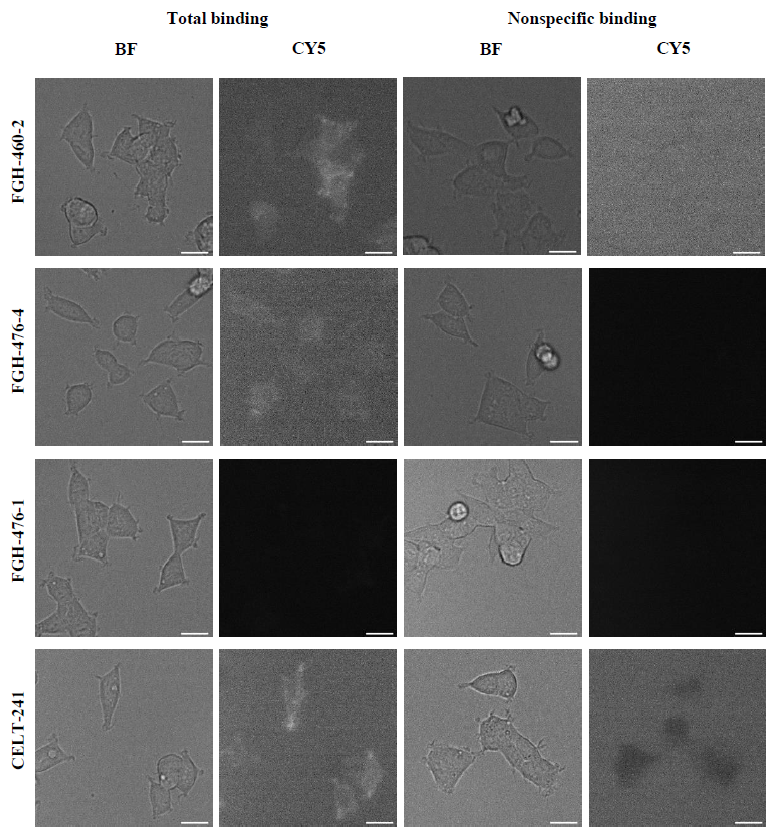

**Supplementary Figure S2**. Ligand binding to the D_2_ receptor. Bright-field (BF) and fluorescence microscopy images were captured 2 h after the start of association. Each well contained HEK293-D2R cells, to which 10 nM fluorescent ligands were added. To determine non-specific binding, 10 μM spiperone solution was added to the wells in addition to the fluorescent ligand solution. Fluorescence intensity was measured in the RFP channel (A), Texas Red (B), or CY5 (C). The scale bar corresponds to 20 μm. For better visualization, the displayed image's intensity range was set according to the minimum and maximum pixel values, using Fiji software's default visualization settings.

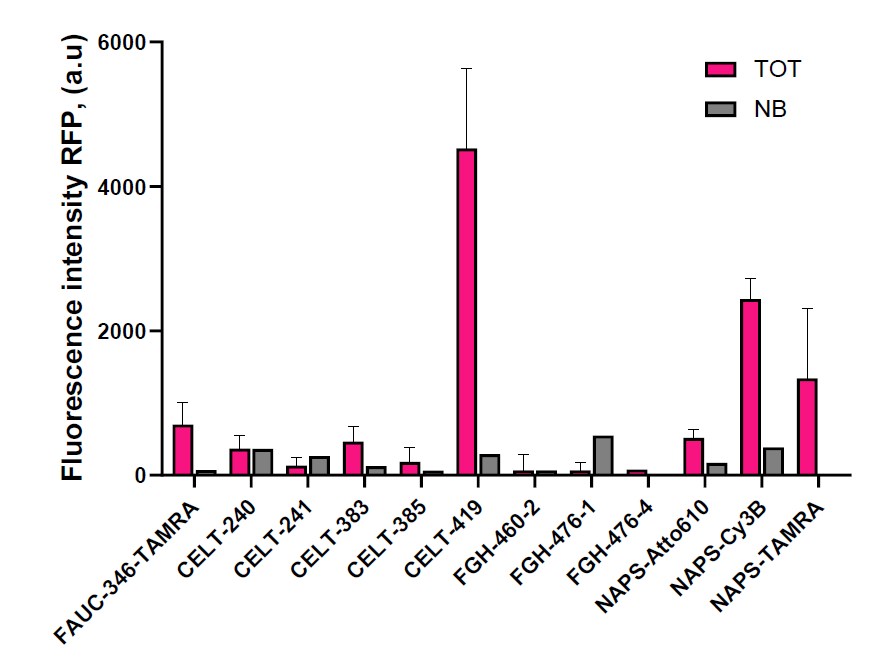

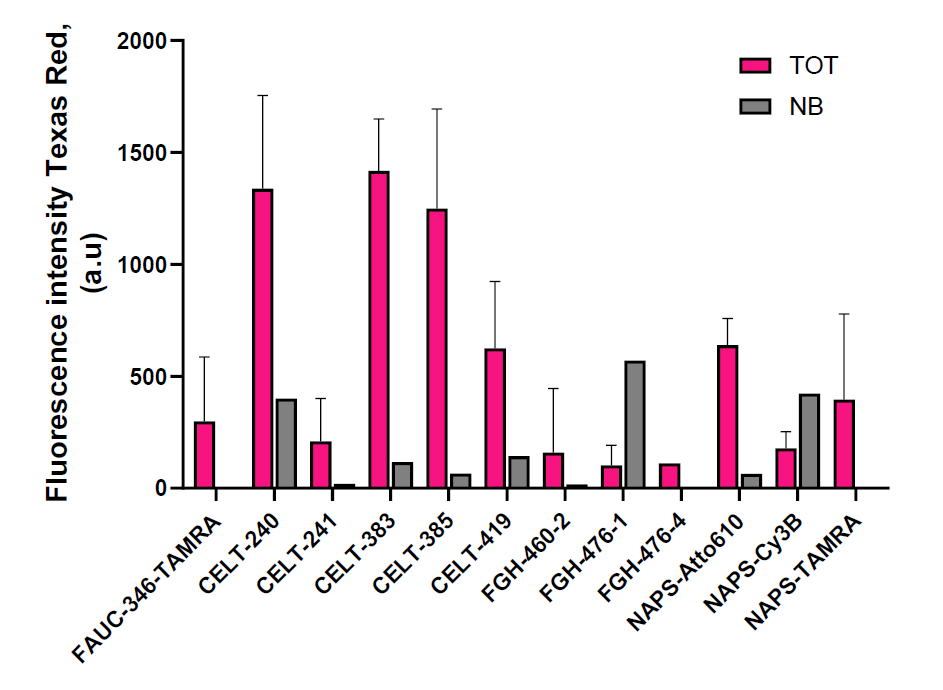

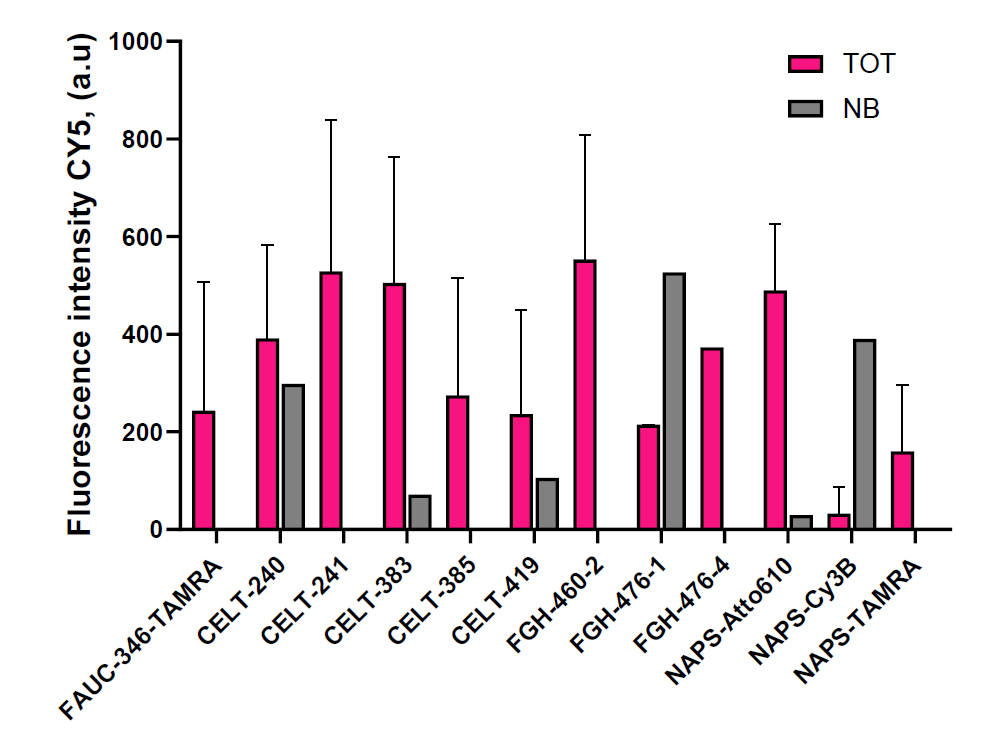

**Supplementary Figure S3**. Quantified fluorescence intensity localized to the HEK293-D2R cells for all tested fluorescent ligands, in the RFP (A), Texas Red (B) and CY5 (C) channels. Total (TOT) and non-specific (NB) binding are presented as means with standard deviation (SD).

**Supplementary Table S1.** Signal-to-noise ratios of fluorescent ligands in different fluorescence channels.

| Fluorescent ligand | Z' (CY5 channel) | Z' (RFP channel) | Z' (TEXAS RED channel) |
| --- | --- | --- | --- |
| FAUC-346-TAMRA | -2.18 | -0.42 | -1.85 |
| CELT-240 | -5.14 | -430.95 | -0.33 |
| CELT-241 | 0.08 | -1.70 | -2.01 |
| CELT-383 | -0.78 | -0.85 | 0.47 |
| CELT-385 | -1.06 | -3.95 | -0.13 |
| CELT-419 | -3.91 | 0.22 | -0.86 |
| FGH-460-2 | -0.29 | -236.9 | -5.04 |
| FGH-476-1 | 1.00 | 0.37 | 0.43 |
| FGH-476-4 | n/a | n/a | n/a |
| NAPS-Atto610 | 0.11 | -0.04 | 0.38 |
| NAPS-Cy3B | 0.55 | 0.58 | 0.07 |
| NAPS-TAMRA | -0.37 | -1.15 | -1.15 |
